# Nascent peptide-induced translation discontinuation in eukaryotes impacts biased amino acid usage in proteomes

**DOI:** 10.1101/2022.01.27.477990

**Authors:** Yosuke Ito, Yuhei Chadani, Tatsuya Niwa, Ayako Yamakawa, Kodai Machida, Hiroaki Imataka, Hideki Taguchi

**Affiliations:** School of Life Science and Technology, Tokyo Institute of Technology, Yokohama 226-8503, Japan; Cell Biology Center, Institute of Innovative Research, Tokyo Institute of Technology, Yokohama 226-8503, Japan; Graduate School of Engineering, University of Hyogo, Himeji, Hyogo 671-2280, Japan

**Author notes:** These authors contributed equally to this work (Yosuke Ito, Yuhei Chadani).

**Keywords:** ribosome, translation, nascent polypeptide, ribosome tunnel, *Saccharomyces cerevisiae*, translation abortion, peptidyl-tRNA hydrolase, amino acid repeat

## Abstract

Robust translation elongation of any given amino acid sequence is a prerequisite to shape proteomes. Nevertheless, nascent peptides could destabilize ribosomes, since consecutive negatively charged residues in bacterial nascent chains stochastically can induce discontinuation of translation, in a phenomenon termed intrinsic ribosome destabilization (IRD). Here, we show that IRD also occurs in eukaryotic translation. Nascent chains enriched in aspartic acid (D) or glutamic acid (E) in the N-terminal regions could prematurely terminate translation, producing premature products as peptidyl-tRNA species. Although eukaryotic ribosomes are more robust to ensure uninterrupted translation, we found many endogenous D/E-rich peptidyl-tRNAs in the N-terminal regions in cells lacking a peptidyl-tRNA hydrolase, indicating that the translation of the N-terminal D/E-rich sequences poses an inherent risk. Indeed, a bioinformatics analysis revealed that the N-terminal regions of ORFs avoid D/E enrichment, implying that the translation defect partly restricts the overall amino acid usage in proteomes.

## Introduction

Life depends on the functions of proteins. Myriads of proteins in living organisms are synthesized by ribosomes according to translation mechanisms. Robust translation elongation of any given amino acid sequence is a prerequisite to shape proteomes. The translation mechanisms of ribosomes, including the essential properties of the peptidyl transferase center (PTC) and the exit tunnel, are well conserved in all living organisms. The ribosome complex processively synthesizes the nascent polypeptide chain without detachment, despite the large conformational changes during the elongation cycle. The elongation by the translating complex progresses on the open reading frame (ORF) of an mRNA until the ribosome arrives at a stop codon. Essentially, the ribosomes decode any codon on ORFs. The growing nascent polypeptides pass through the ribosomal exit tunnel (Wilson & Beckmann, 2011), which encloses 30~40 amino acids of the growing nascent polypeptidyl-tRNA(Dao Duc *et al*, 2019; Ito & Chiba, 2013).

Recent studies have revealed that the translation elongation rhythm is non-constant (Ito & Chiba, 2013). The dynamics of translating ribosomes is affected not only by the scarcity of codons or tRNAs and mRNA secondary structures (Yanofsky, 1981; Wolin & Walter, 1988; Wen *et al*, 2008; Choi *et al*, 2015) but also by specific interactions between the ribosome exit tunnel and the nascent chain (Ito & Chiba, 2013; Chadani *et al*, 2016; Buskirk & Green, 2017). The tunnel-nascent chain interactions result in altered translational speeds in both bacteria and eukaryotes (Bhushan *et al*, 2010; Seidelt *et al*, 2009; Bhushan *et al*, 2011; Shanmuganathan *et al*, 2019). Mechanistically, the electrostatic interaction between negatively charged ribosomal RNAs and positively charged nascent polypeptides could cause ribosomal pausing (Lu & Deutsch, 2008; Charneski & Hurst, 2013; Tuller *et al*, 2011). The translation of proline stretches is known to cause elongation pausing, which is alleviated by eIF5A (Gutierrez *et al*, 2013). Recent global studies on eIF5A using the ribosome profiling method showed that eIF5A also affects the elongation of tripeptide motifs enriched with negatively charged amino acids (Asp: D and Glu: E), in addition to proline stretches (Schuller *et al*, 2017; Pelechano & Alepuz, 2017). The translation of aberrant mRNAs causes elongation pausing and ribosome collision, and is subsequently eliminated by ribosome-associated quality control (RQC) systems in eukaryotes (Brandman *et al*, 2012; Inada, 2017).

In addition to translation pausing, nascent chains even destabilize the translating ribosome from within. We recently found that *Escherichia coli* ribosomes are destabilized by a class of amino acid sequences in nascent chains, typically containing D/E-runs such as ten consecutive glutamates (10E), stochastically leading to a translation termination termed intrinsic ribosome destabilization (IRD, Chadani *et al*., 2017, 2021). The peptidyl-tRNAs produced by IRD are cleaved by peptidyl-tRNA hydrolase (Pth). Pth, an essential enzyme in bacteria, recycles the peptidyl-tRNAs outside the ribosome complex by cleaving the ester bonds between the peptides and tRNAs. Although the translation of negatively charged sequences is risky, there are several counteracting mechanisms to prevent premature termination. A high concentration of magnesium, which is known to stabilize the ribosome complex, counteracts IRD. Interestingly, this Mg^2+^ dependency of IRD provides a physiological impact. In addition, most potential IRD sequences in the middle of ORFs remain cryptic by the tunnel-spanning nascent polypeptide preceding the IRD sequences (Chadani *et al*, 2021).

Since translation mechanisms are conserved in all domains of life, one might ask whether IRD takes place even in eukaryotic cells. Compared to bacterial ORFs, eukaryotic ORFs contain many consecutive D/E sequences, which are considered to extend the protein function in eukaryotic organelles such as the nucleus. For example, essential genes including more than 20 D/E runs are present in budding yeast. Here, we show that the premature translation termination induced by nascent chains enriched in D or E residues around N-terminal regions also occurs in eukaryotes, as revealed by an *in vivo* reporter assay and a reconstituted cell-free translation system. The accumulation of the resultant abortive peptidyl-tRNAs is toxic, since yeast cells lacking Pth cannot grow when IRD-prone sequences are overexpressed. We identified many endogenous IRD-derived peptides by a mass spectrometry (MS) analysis of the RNA fractions in the Pth-deletion strain. A bioinformatics analysis unveiled that proteomes have a bias in their amino acid distribution to avoid the premature termination risk induced by IRD. Thus, the translation elongation process disfavors a subset of amino acid sequences harboring D/E-rich residues at their N-terminal regions.

## Results

### Premature termination during the translation of consecutive negatively charged amino acids in the N-terminal regions

We recently found a noncanonical translation dynamics termed IRD, where nascent chains enriched in D/E residues destabilize the translating *E. coli* ribosome and stochastically terminate translation in a context-dependent manner (Chadani *et al*., 2017, 2021). To determine whether noncanonical translation also takes place in eukaryotic cells, we constructed a reporter system based on a dual-luciferase assay to monitor the potential IRD in *Saccharomyces cerevisiae* (**Fig. 1A**). Firefly luciferase (Fluc) and *Renilla* luciferase (Rluc, as an internal expression control) were expressed under the galactose-inducible bidirectional promoter. Fluc was fused to GFP, a homo-decamer of amino acids (10X; 10D, 10E, 10N, 10Q, and 10T), a linker peptide, and a self-cleaving peptide, T2A (**Fig. 1A**). If the translation of the consecutive D/Es such as 10D induced translation abortion, then the ratio of the activity of Fluc to that of Rluc, defined as the translation continuation (TC) index, would decrease.

**Figure 1.**
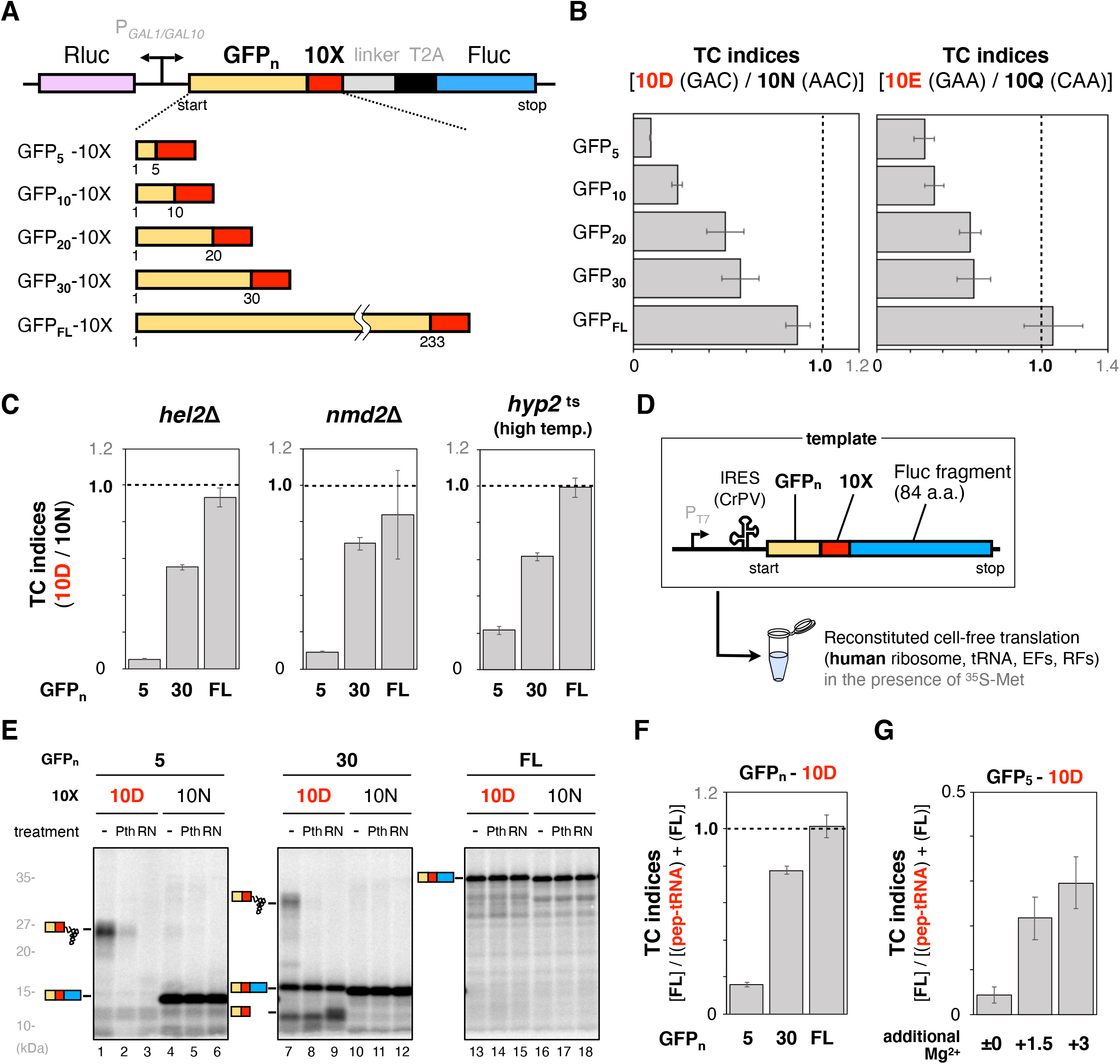
Consecutive negatively charged amino acids in the N-terminal regions prematurely terminate translation. (**A**) Schematic representation of the dual luciferase assay. Homo-decamers of amino acids (10X) were inserted between varying lengths of the N-terminal region of GFP (GFP_*n*_; *n*=5, 10, 20, 30 and 233 as full length: FL) and a linker. Firefly (Fluc) and *Renilla* (Rluc) luciferases were used as a reporter enzyme to investigate the translation continuation and an internal translation control, respectively. The *GAL1-GAL10* promoter is a galactose-inducible bidirectional promoter. T2A is the self-cleaving peptide from *Thosea asigna* virus 2A (Kim *et al*., 2011). (**B**) The translation efficiencies of ORFs harboring 10X in *S. cerevisiae* were evaluated by the reporter system shown in (A). After induction by galactose, the translation efficiencies were measured as the relative activity of Fluc to that of Rluc. Translation continuation (TC) indices were calculated by the following formula: {Fluc activity/Rluc activity (10D)} / {Fluc activity/Rluc activity (10N)}. Error bars indicate standard deviations from three trials. The codons encoding 10X are indicated in parentheses. (**C**) TC indices evaluated using the reporter system in the *hel2*Δ, *nmd2*Δ and *hyp2* temperature sensitive (*ts*) strains are shown. The *hyp2^ts^* cells were grown at 33 °C instead of 30 °C, the normal cultivation temperature. (**D**) Schematic representation of the fusion gene, GFP_*n*_-Methionine-10X-truncated Fluc, for the cell-free assay. A methionine codon was inserted between GFP and 10X to label the translation product with ^35^S-methionine. (**E**) The fusion genes were translated using the human factor-based reconstituted cell-free translation system (HsPURE system) including ^35^S-methionine. The products were treated with a peptidyl-tRNA hydrolase (yeast Pth2p: *Pth*) as indicated and separated by neutral pH SDS-PAGE with optional RNase A (*RN*) pretreatment. Radioactive bands were detected by a phosphorimager. Peptidyl-tRNAs and tRNA-cleaved polypeptides are indicated by schematic labels. (**F**) TC indices calculated from the band intensities in (E). The TC index was defined by the following formula: {completed chain (10D)} / {completed chain (10D) + peptidyl-tRNA (10D)}. Error bars indicate standard deviations from three trials. (**G**) Effect of Mg^2+^ on the 10D sequence-dependent translation attenuation. The GFP_5_-10D sequences were translated by the HsPURE system supplemented with additional Mg^2+^ as indicated, and individual TC indices were calculated.

The TC indices for the 10D and the 10E fused with full-length GFP (GFP_FL_) were 0.9~1.1, when the indices were normalized by those of 10N and 10Q, respectively, to eliminate the effect of the negative charges in the motifs (**Fig. 1B**), showing uninterrupted translation continuation in these 10D and 10E constructs. Since IRD in *E. coli* tends to occur very early in the coding sequence (Chadani *et al*., 2021), we then shortened the GFP moiety to 5 (GFP_5_) and 30 (GFP_30_) amino acids from the N-terminus. Strikingly, the TC indices decreased depending on the GFP length in both 10D/10N and 10E/10Q: a shorter GFP length led to a lower TC index (**Fig. 1B**). A quantitative MS analysis of peptides derived from the two luciferases also confirmed that the relative luciferase activities were in accordance with the translation products (**Fig. S1A**). The TC indices of 10D/10T, where the 10T sequence was encoded in (ACG)_10_, frame-shifted codons of (GAC)_10_ for the 10D, also depended on the GFP length as in the case of 10D/10N, confirming that the amino acid sequence, but not the RNA sequence, is critical for the reduction (**Fig. S1B**).

The reduced luciferase activities in the 10D and 10E constructs might be due to mRNA quality control systems, including no-go decay (NGD, Matsuo *et al*, 2017; Doma & Parker, 2006) and nonsense-mediated decay (NMD, He & Jacobson, 1995; Cui *et al*, 1995). To address this, we used the *hel2*Δ and *nmd2*Δ strains, with impaired NGD-mediated ribosome dissociation and NMD-mediated clearance of aberrant mRNA, respectively. The 10D-dependent reduced TC indices in the constructs harboring shorter GFP moieties were not affected in both strains (**Fig. 1C**), confirming that NGD and NMD are not responsible for the reduced translation. In addition, the TC indices of GFP_5_-10D (or GFP_30_-10D) were not altered even in a temperature-sensitive strain of *hyp2*, which encodes the eIF5A protein (**Fig. 1C**, (Li *et al*, 2011), after we verified the inefficient translation of 10 consecutive prolines in the *hyp2*^ts^ strain at a semi-lethal temperature, 33 °C, using the dual-luciferase system (**Fig. S1C and S1D**). Taken together, the major translation quality control systems known to date are not involved in the consecutive D/E-dependent translation attenuation.

### D/E-run-dependent translation abortion in a reconstituted cell-free translation system

The possibility of elongation pausing, and the involvement of other *trans*-acting factors for the translation attenuation, cannot be assessed by the above reporter assay. Therefore, we adopted a human factor-based reconstituted cell-free translation system, the HsPURE system (Machida *et al*, 2014), to ultimately determine whether the decreased translation is intrinsically induced by the nascent peptide chain. We constructed fusion genes encoding GFP (GFP_FL_, GFP_5_, or GFP_30_)-10X-Fluc (N-terminal 84 a.a. fragment) and translated them with the HsPURE system (**Fig. 1D**). The translated products were separated by neutral pH SDS-PAGE, which can analyze intact peptidyl-tRNA species (Chadani *et al*, 2016, 2017, 2021). All of the GFP_FL_-containing constructs produced full-length polypeptides, regardless of whether 10X was 10D or 10N (**Fig. 1E**, *lanes 13*, *16*). In the constructs harboring GFP_30_, the 10D construct produced several bands (*lane 7*), whereas the major band from the 10N construct was the full-length polypeptide, which is insensitive to RNase (*lane 10*). The largest band in GFP_30_-10D (*lane 7*) was sensitive to RNase (*lane 9*), indicating that the band is a peptidyl-tRNA. In addition, we found that one of the Pths in *S. cerevisiae* (Pth2: the reason why Pth2 was chosen is shown in **Fig. 3**) cleaved the resultant peptidyl-tRNA (*lane 8*), suggesting that the peptidyl-tRNAs are derived from the destabilization of the ribosome complex, rather than translational pausing (Chadani *et al*, 2017, 2021) In the construct harboring the shorter GFP sequence (GFP_5_), the translation produced more RNase- and Pth-sensitive peptidyl-tRNA species (*lane 1*). The TC indices calculated from the quantification of the band intensities in the HsPURE system (**Fig. 1F**) were consistent with those obtained from the luciferase reporter assay in yeast (**Fig. 1B**).

**Figure 2.**
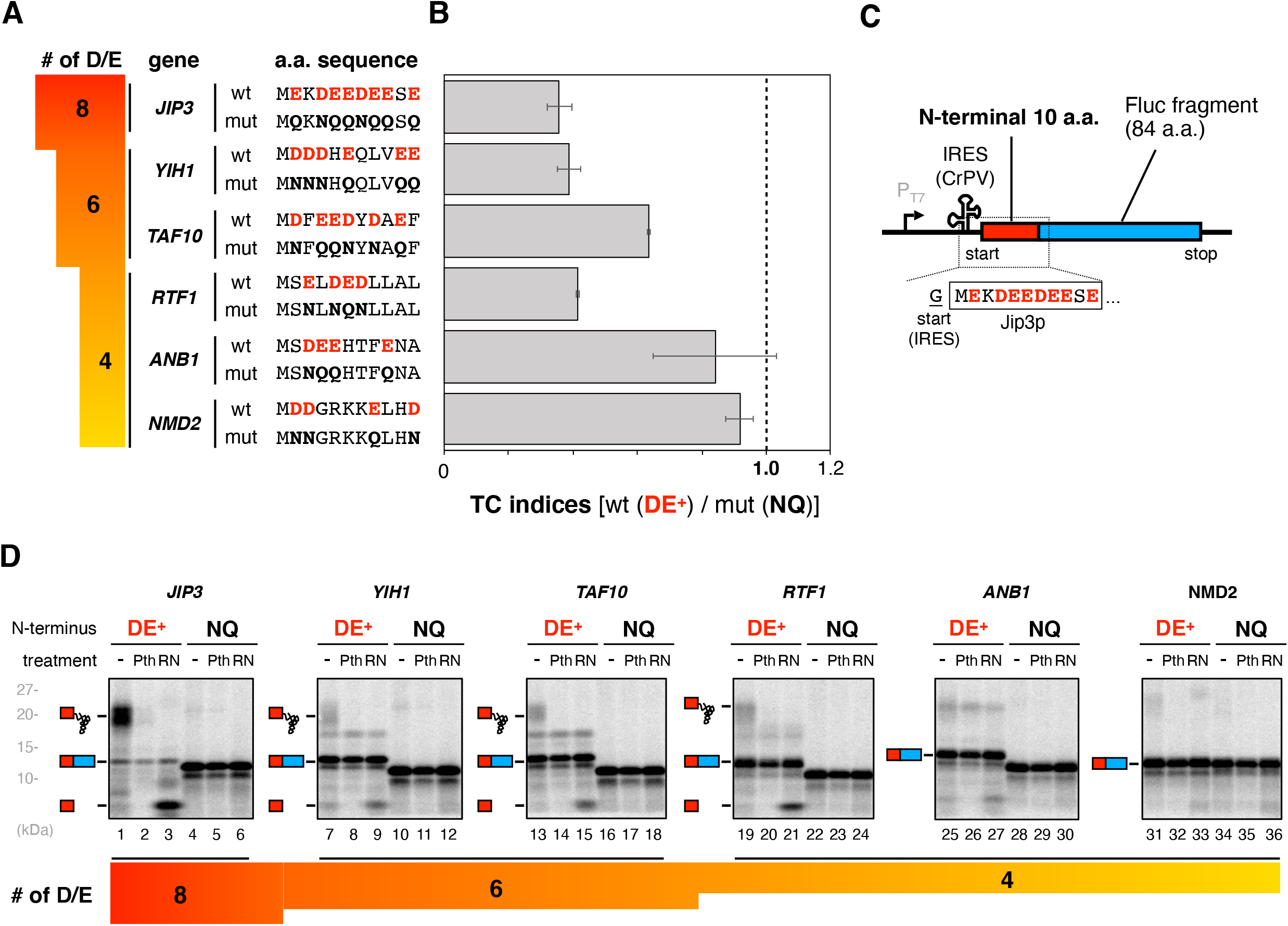
Translation of endogenous genes that have multiple Asp/Glu (D/E) residues in the N-terminal regions. (**A**) The list of representative genes with 8, 6, and 4 D/Es in the N-terminal 11 amino acids. The N-terminal sequences (*wt*) are shown with the mutant sequences (*mut*), in which D/Es are substituted by N/Qs. (**B**) The N-terminal 10 amino acid sequences except for the first methionine were inserted instead of the 10X in the dual luciferase reporter construct shown in Fig. 1A. TC indices were calculated by the following formula: {Fluc activity (*wt*) / Rluc activity (*wt*)} / {Fluc activity (*mut*)/Rluc activity (*mut*)}. Error bars indicate standard deviations from three trials. (**C and D**) The N-terminal 11 amino acids of the genes with multiple D/Es were fused with truncated Fluc (**C**). (**D**) The genes were translated using the HsPURE system, as described in Fig. 1E.

**Figure 3.**
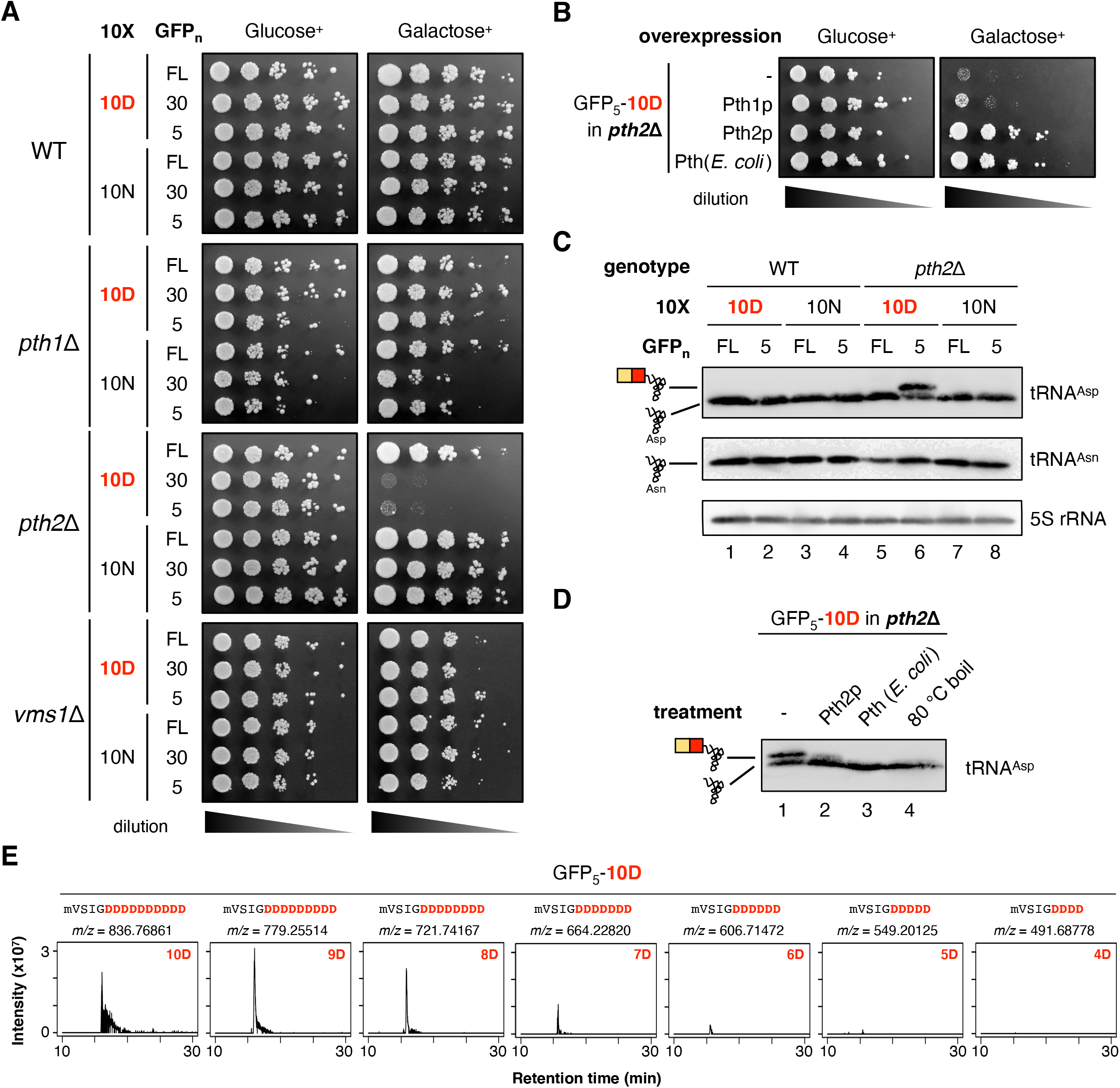
Pth2, one of the peptidyl-tRNA hydrolases, is responsible for hydrolyzing the peptidyl-tRNAs derived from the D/E-rich sequence-induced premature termination. (**A**) Wild type (*WT*), *pth1*Δ, *pth2*Δ and *vms1*Δ strains harboring the plasmids for the expression of the 10X-containing ORFs shown in Fig. 1A (the 10D or 10N series) were spotted on SD (Glucose^+^) or SG (Galactose^+^) plates lacking uracil and cultivated at 30 °C for 2 days. Cultures were 10-fold serially diluted in distilled water before spotting. (**B**) Complementation assay of Pth variants. The *pth2*Δ mutant harboring the GFP_5_-10D construct showing the lethal phenotype, the transformed empty vector and plasmids carrying *pth1*, *pth2*, and *pth* from *E. coli*, were spotted on SD or SG plates lacking uracil and leucine and incubated at 30°C for 2 days. (**C**) Northern blot analysis with anti-tRNA^Asp^ and tRNA^Asn^ probes showing tRNA^Asp^ with or without a short peptide (peptidyl-tRNA or tRNA, respectively) depending on strains and constructs. The results of 5S rRNA are also shown as a loading control. (**D**) Northern blot analysis with the anti-tRNA^Asp^ probe, showing tRNA^Asp^ with or without a short peptide depending on the hydrolysis treatment. Pth2: RNA extract with 1 μM purified Pth2 was incubated at 30 °C for 20 min. Pth (*E. coli)* RNA extract with 1 μM purified *E. coli* Pth was incubated at 37 °C for 20 min. 80 °C 20 min (pH 11.0): RNA extract was boiled under alkaline conditions (Ito *et al.*, 2011). (**E**) Extracted ion chromatograms of the peptide fragment expressed from the GFP_5_-10D sequence in yeast *pth2*Δ cells. Each panel represents an extracted ion chromatogram of MS1 for a monoisotopic ion of a GFP_5_-(X)D peptide with a specific m/z value, as indicated at the top. The letter “m” in the sequence means oxidized methionine.

We next evaluated the Mg^2+^ concentration dependency of the translation abortion since IRD in *E. coli* is suppressed by high Mg^2+^ concentrations, which stabilize the ribosome subunit association (Schlessinger *et al*, 1967; Ron *et al*, 1968; Chadani *et al*, 2017). Additional Mg^2+^ in the cell-free translation system alleviated the premature translation termination in GFP_5_-10D (**Fig. 1G**) and GFP_30_-10D (**Fig. S1E**), but did not affect the translational pausing in hCMV uORF2 (**Fig. S1F)**, suggesting that the premature termination is somehow associated with destabilization of the ribosome, similar to that in *E. coli*.

These *in vivo* and *in vitro* results demonstrated that the translation of the N-terminal D/E-runs in eukaryotes also destabilizes the translating ribosomes, without requiring any factor other than the essential translation components, eventually resulting in the accumulation of immaturely terminated products.

### Endogenous ORFs harboring D/E-rich sequences in the N-terminal regions induce premature termination

We next examined yeast ORFs harboring multiple D/E residues in the N-terminal regions to determine whether such D/E-rich sequences in endogenous ORFs induce premature termination. We found 169 ORFs in *S. cerevisiae* that contain 4 or more D/E residues in the first 10 residues after the initial methionine. One ORF (*JIP3*) with 8 D/Es was the maximum (**Fig. 2A**). In addition to this ORF with the 8 D/Es, we selected two and three ORFs with 6 and 4 D/Es, respectively, in the N-terminal regions to investigate whether IRD occurs during the translation of these candidates (**Fig. 2A**). The dual-luciferase reporter assay revealed that more than 60% of the translation of the ORF with 8 D/Es was discontinued immaturely (**Fig. 2B**). We also observed a relatively high rate of discontinuation during the translation of the two ORFs with 6 D/Es. In contrast, only one ORF with 4 D/Es (*RTF1*) exhibited high discontinuation efficiency (**Fig. 2B**).

We also performed the HsPURE *in vitro* translation to evaluate the accumulation of abortive peptidyl-tRNAs for these six endogenous ORFs (**Fig. 2C**). The cell-free translation of *JIP3* harboring 8 D/Es in the N-terminal region preferentially produced RNase- and Pth-sensitive peptidyl-tRNA species (**Fig. 2D**). We also observed the accumulation of such peptidyl-tRNAs during the translations of two ORFs with 6 D/Es and one ORF with 4 D/Es (*RTF1*), albeit to lesser extents as compared to the ORF with 8 D/Es (**Fig. 2D**). The translation reactions of other ORFs with 4D/Es (*ANB1*, *NMD2*) hardly accumulated any band that was sensitive to RNase or Pth (**Fig. 2D**), consistent with the corresponding data obtained by the dual-luciferase reporter assay (**Fig. 2B**). Collectively, these results revealed that the endogenous ORFs harboring the D/E-rich sequences at the N-terminal regions also induce the translation discontinuation, which is roughly proportional to the number of D/Es in the regions, as shown in the previous study (Chadani *et al*., 2017). In addition, the higher Mg^2+^ concentration abrogated the accumulation of peptidyl-tRNA in the 8 D/E-containing JIP3p (**Fig. S2**), further supporting the idea that IRD is involved in the accumulation of the peptidyl-tRNA, leading to the translation abortion during the translation of endogenous ORFs.

The above results demonstrated that the N-terminal D/E-rich sequences abort translation. In contrast, when the D/E-runs were translated after GFP_FL_, most of the IRD was circumvented even in the translation of GFP_FL_-20D (or −20E) (**Fig. S2B**). Since the *E. coli* ribosome was unable to translate GFP_FL_-20D (or −20E) due to IRD (**Fig. S2B**), the eukaryotic ribosome is more robust to continue translation elongation.

### D/E-rich residues-induced peptidyl-tRNAs are cleaved by Pth2 *in vivo*

The peptidyl-tRNA species produced by translation abortion would be cleaved by a Pth (peptidyl-tRNA hydrolase). In *E. coli*, the accumulation of peptidyl-tRNAs generated by premature translation termination is toxic, mainly due to the scarcity of tRNAs (Menninger, 1979; Heurgué-Hamard *et al*, 1996). So far, *S. cerevisiae* has two known Pths, encoded by *PTH1* and *PTH2* (Rosas-Sandoval *et al*, 2002; Menez *et al*, 2002). In addition, recent studies revealed that the 60S-accomodated peptidyl-tRNA is cleaved by Vms1p, which removes the CCA end of tRNA (Verma *et al*, 2018; Rendón *et al*, 2018; Kuroha *et al*, 2018). If one of the Pths is involved in the cleavage of the peptidyl-tRNAs, then the accumulation of undigested peptidyl-tRNAs in a strain lacking the responsible Pth gene would be potentially toxic.

Accordingly, we examined whether the overexpression of the D/E-runs affected the growth of the *pth* deletion strains. After we confirmed that the three single deletion strains, *pth1*Δ, *pth2*Δ, and *vms1*Δ, were as viable as the wild-type strain, fusion genes containing GFP-10X, where GFP sequences (GFP_FL_, GFP_5_ or GFP_30_) are followed by 10X (10D or 10N), were expressed under a galactose-inducible promoter (**Fig. 3A**). Expression of the genes in all combinations did not affect the viabilities of the *pth1*Δ and *vms1*Δ strains, and the wild-type (**Fig. 3A**). In contrast, the expression of genes with truncated GFPs followed by 10D (GFP_5_-10D and GFP_30_-10D), but not those by the 10N or 10T series, in the *pth2*Δ strain was lethal (**Fig. 3A, Fig. S3A**). The expression of the GFP_5_- or GFP_30_-10E instead of that of the 10D counterparts also exhibited toxicity (**Fig. S3B**), indicating that the D/E-runs are critical for the harmful phenotype. The lethal phenotype of *pth2*Δ upon the expression of GFP_5_-10D was rescued by the expression of Pth2p or *E. coli* Pth, but not Pth1p (**Fig. 3B**).

The results of this phenotype assay strongly support the assumption that the accumulation of the IRD-derived peptidyl-tRNAs in the *pth2*Δ strain is toxic. To obtain direct evidence for the accumulation of peptidyl-tRNAs in the *pth2*Δ strains, we conducted a Northern blot analysis to detect the peptidyl-tRNA species. Total RNA fractions were individually isolated from wildtype and *pth2*Δ strains by phenol extraction, and then electrophoresed, electroblotted, and probed by an anti-tRNA^Asp^ oligonucleotide. We detected a higher molecular weight band, in addition to the tRNA^Asp^ band, in the total RNA isolated from the *pth2*Δ strain with the GFP_5_-10D overexpression (**Fig. 3C**). This higher molecular weight band was not observed in the Northern blot for the GFP_5_-10N using the tRNA^Asp^ oligonucleotide, confirming that the consecutive Asp residues were responsible for the higher molecular weight band. The higher molecular weight band in the *pth2*Δ lysate was sensitive to treatments with Pth proteins, Pth2p or *E. coli* Pth, or heat under high pH conditions, further supporting that this band is the abortive product of the GFP_5_-10D translation (**Fig. 3D**).

Final proof for the accumulation of the peptidyl-tRNAs in the *pth2*Δ strain was provided by the MS analysis. The MS analysis of the *pth2*Δ lysate expressing GFP_5_-10D identified several unique peptides derived from the peptidyl-tRNAs: MS signals for GFP_5_-6D, 7D, 8D, 9D and 10D in the GFP_5_-10D sequence (**Fig. 3E**, **Fig. S3C, D**). Such stochastic production of various numbers of negatively charged residues in the peptidyl-tRNAs also occurs in IRD in *E. coli* (Chadani *et al*., 2017).

Taken together, we concluded that the nascent negative-charge amino acid stretches at the N-terminal regions induce IRD, stochastically terminate translation, and eventually produce peptidyl-tRNAs in the eukaryotic translation system. In the wild-type yeast strain, the peptidyl-tRNAs are cleaved by Pth2p to prevent the harmful accumulation of the abortive products. Accordingly, the Pth2 protein would be essential for cell viability under stress conditions, where the abortive peptidyl-tRNAs are highly accumulated.

### Accumulation of endogenous peptidyl-tRNAs with D/E-enriched C-termini in *pth2*Δ cells

A series of experiments in the *pth2*Δ strain prompted us to broadly explore the endogenous ORFs that prematurely terminate the translation in an IRD-dependent manner. To this end, we developed a new MS method to analyze endogenously accumulated abortive peptidyl-tRNAs in the *pth2*Δ lysate. The method is a combination of the isolation of the RNA fraction containing peptidyl-tRNAs (Chomczynski & Sacchi, 1987), the heat- and high pH-induced cleavages of the peptide moieties from the peptidyl-tRNAs (Ito *et al*, 2011), and the identification of the resultant peptides by MS-based shotgun proteomics (**Fig. 4A**).

**Figure 4.**
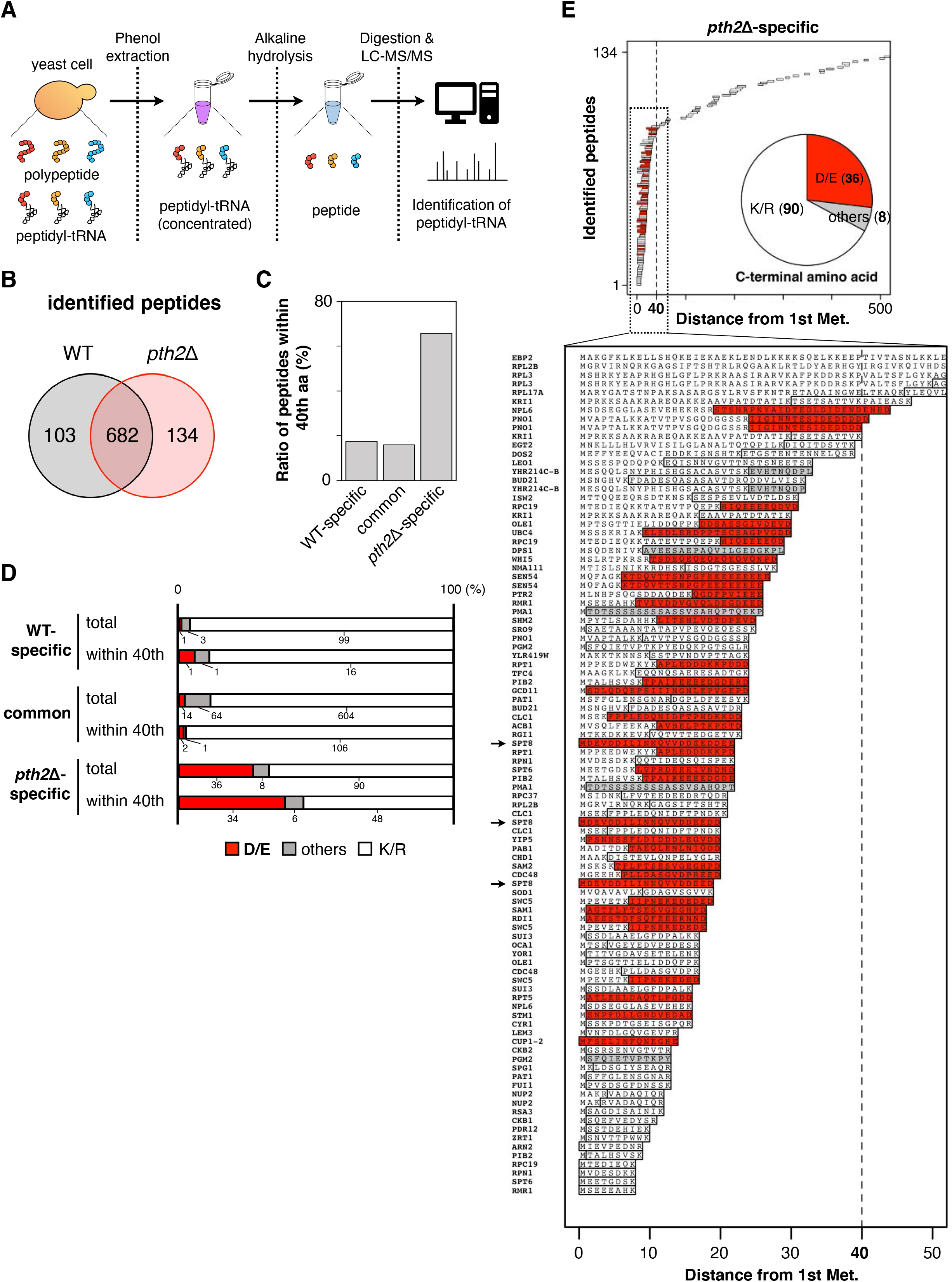
Identification of the abortive peptidyl-tRNAs accumulated in the *pth2*Δ strain by LC-MS/MS analysis. (**A**) Flowchart of the analysis of the abortive peptidyl-tRNAs in lysates, using LC-MS/MS. Peptidyl-tRNAs, which are enriched in the phenol-extracted RNA fraction from wild type (WT) or *pth2*Δ strains, are chemically hydrolyzed and then digested by trypsin and LysC peptidase. The resultant peptides are identified by LC-MS/MS. Experimental details are described in the Methods section. (**B**) The numbers of peptides identified in each strain are represented in the Venn diagram. (**C**) The percentages of the number of detected peptides whose C-terminal positions were within 40 amino acids from the first methionine. (**D**) The C-terminal amino acids of identified peptides detected in WT, common or *pth2*Δ, whose C-terminal positions were within 40 amino acids from the first methionine. The stacked bar graphs represent the C-terminal amino acid of identified peptides (red: D/E; gray: others; white: K/R). (**E**) Mapping of identified peptides specific to the *pth2*Δ strain. Map charts representing the localization of identified peptides on each protein, with the enlarged view presented below. The dotted line indicates the position of 40 amino acids from the first methionine. The length of the 40 amino acids is roughly equivalent to that of the exit tunnel in the ribosome. *Insets*: Pie chart representing the amino acids at the C-termini of identified peptides (same as Fig. 4D, red: D/E; gray: others; white: K/R).

We identified 816 peptides from the RNA fraction in the *pth2*Δ lysate (**Fig. 4B**). Among them, 682 overlapped with the peptide set in the wild-type lysate, whereas 134 peptides were specific to those in the *pth2*Δ cells (**Fig. 4B**). The *pth2*Δ-specific peptides derived from the peptidyl-tRNAs have notable distinguishing features. First, the *pth2 Δ-specific* peptides were enriched at the N-terminal regions of the ORFs, as compared to the wild type-specific or common peptides: 65% (88 out of 134) of the identified peptides were mapped within the 40 amino acids from the start methionine in the annotated ORFs (**Fig. 4C**). Since the ~40 amino acid residues are considered to be accommodated in the ribosome exit tunnel, the enrichment suggests that the *pth2*Δ-specific peptides are produced by premature termination before the N-terminal portions fully occupy the tunnel. Second, Asp (D) or Glu (E) were enriched in the C-terminal ends of the *pth2*Δ-specific peptides (**Fig. 4D**). Although the conventional C-terminal residues in the shotgun proteomics analyses of trypsin-treated proteins are Lys (K) or Arg (R) due to the enzymatic properties of trypsin, we used a mode to identify the peptides independently from the C-terminal K/R. The analysis revealed that ~27% (36 out of 134) of the *pth2*Δ-specific peptides had D/Es at the C-termini (**Fig. 4D**). Notably, the preference of D/Es at the C-termini was more pronounced (~39%) if we limited the *pth2*Δ-specific peptides to those mapped within the 40 amino acids from the start methionine (**Fig. 4D and 4E**). Such enrichment of the C-terminal D/Es was only observed in the subset of *pth2*Δ-specific peptides (**Fig. 4D and 4E, Fig. S4A, B**), suggesting that the peptides with the C-terminal D/Es were not derived from trypsin miscleavage, but from the premature termination induced by IRD. One representative ORF meeting these features is *SPT8*. We identified three peptides from the N-terminus of Spt8p, 1-MDEVDDILINNQVVDDEED-19, 1-MDEVDDILINNQVVDDEEDD-20, and 1-MDEVDDILINNQVVDDEEDDEE-22, reflecting the stochastic nature of the termination within the D/E-rich peptide (**Fig. 4E**). Taken together, the peptidyl-tRNA analysis developed here revealed that the 36 D/E-rich peptides from the N-terminal regions of the 27 endogenous ORFs were enriched in the *pth2*Δ cells.

To verify that the premature termination actually occurs during the translation of the genes whose peptidyl-tRNAs were detected in the *pth2*Δ strain, we examined candidate ORFs *in vitro* and *in vivo* (**Fig. 5, Fig. S5**). Translations of seven ORFs using the HsPURE system resulted in the D/E-dependent accumulation of Pth-sensitive peptidyl-tRNA species (**Fig. 5B, Fig. S5B**), confirming that the premature termination of these ORFs indeed occurs in the reconstituted cell-free translation. In addition, the overexpression of three ORFs (*PIB2, SPT8*, and *SEN54*) in the *pth2*Δ strain induced lethal phenotypes in a D/E residue-dependent manner (**Fig. 5C**), probably reflecting the detrimental effect of the abortive peptidyl-tRNAs on the cell viability. These results further confirmed that the translation of the N-terminal D/E-rich sequences in the endogenous mRNAs could cause premature translation termination.

**Figure 5.**
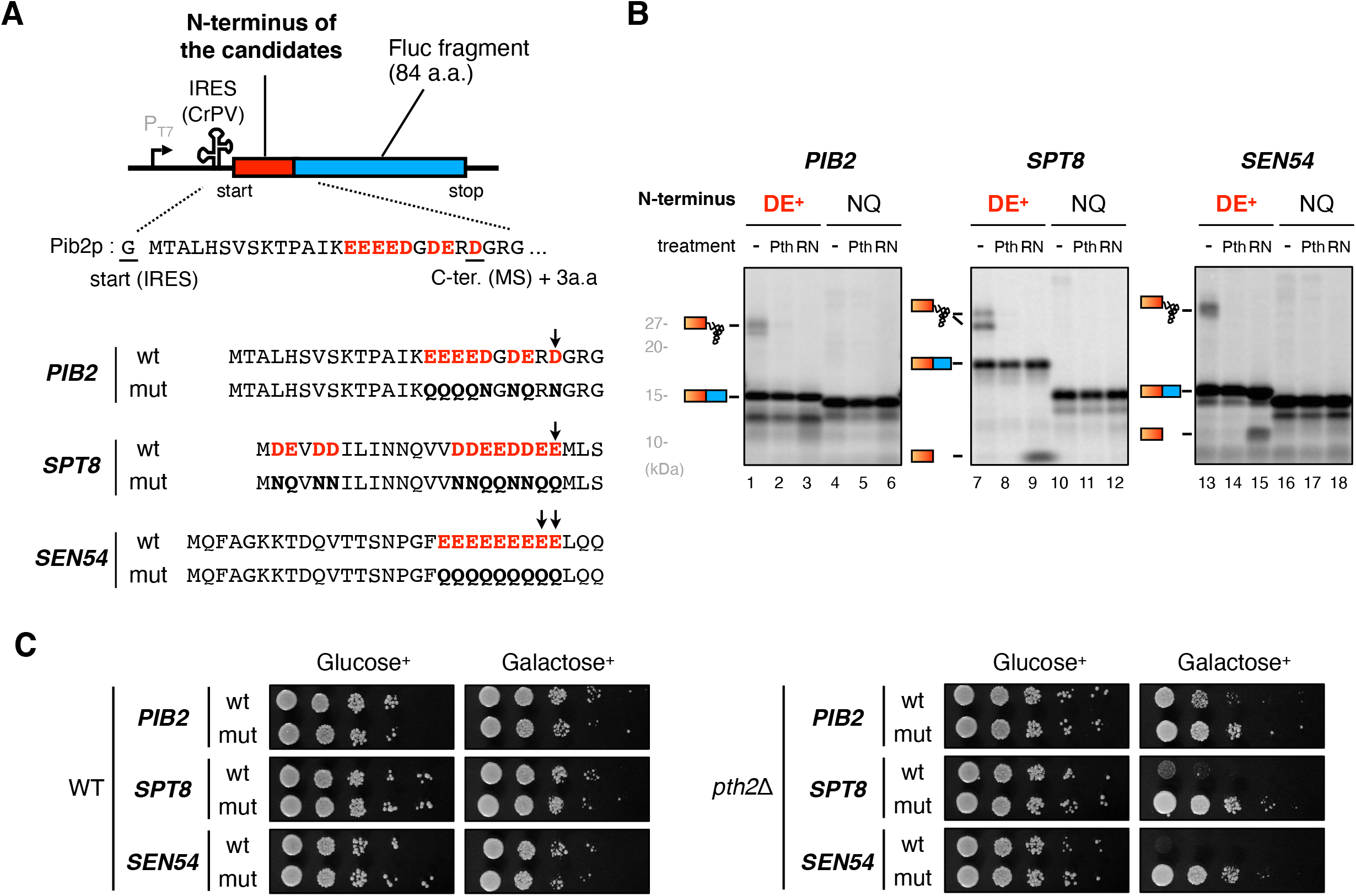
IRD during the translation of LC-MS/MS-identified endogenous genes. (**A**) Schematic representation of the sequences from endogenous genes for the HsPURE *in vitro* translation assay and the spot assay. N-terminal amino acid motifs detected by MS in *pth2*Δ cells (with 3 a.a. extension) were substituted for GFP_n_-10X of the constructs in Fig. 1A and Fig. 1D. (**B**) IRD during the translation of endogenous genes depending on the D/E-rich sequences in the vicinity of the N-terminal region. The reporter genes shown in Fig. 5A were translated using the HsPURE system, as described above. (**C**) Toxicity upon the expression of endogenous IRD-inducing motifs. Wild-type (WT) and *pth2*Δ strains harboring plasmids expressing Fluc fused with the N-terminus of the candidates were spotted on SD and SG plates lacking uracil and grown at 30 °C for 2-3 days.

### Proteomes avoid negatively charged amino acid clusters in the N-terminal regions

The above results implied that the translation of D/E-rich amino acids in the N-terminal regions poses an inherent risk of premature translation termination, from bacteria to human. Accordingly, we assumed that during evolution, organisms would have avoided the translation of D/E-rich amino acids occurring very early in the coding sequences. Therefore, we conducted a bioinformatics analysis to investigate the distribution of the D/E-rich sequences in proteomes.

We extracted 100 amino acids from the N-terminal, middle and C-terminal regions of all known yeast ORFs (5,540 ORFs) and then counted the number of proteins with three or more D/E residues in each ten-residue moving window (**Fig. 6A**). Strikingly, we found that the number of proteins with ≥3 D/E residues in the N-terminal regions dropped early, from the N-terminus to 30-40 amino acids, which is similar to the length of the tunnel. The N-terminal regions are critical in this trend, since there was no such reduction in the middle or C-terminal regions (**Fig. 6A**). The trend for the disfavored D/E usage was i) specific to the D/E residues since there is no such decline in either N/Q or K/R residues (**Fig. 6B** and **Fig. S6A**), ii) statistically significant (**Fig. S6B**), iii) more obvious in the analysis with ≥5 D/E residues (**Fig. S6C**), and iv) not due to a bias in nucleic acid sequences, since there is no such reduction in the frameshifted DNA sequences (**Fig. S6D**).

**Figure 6.**
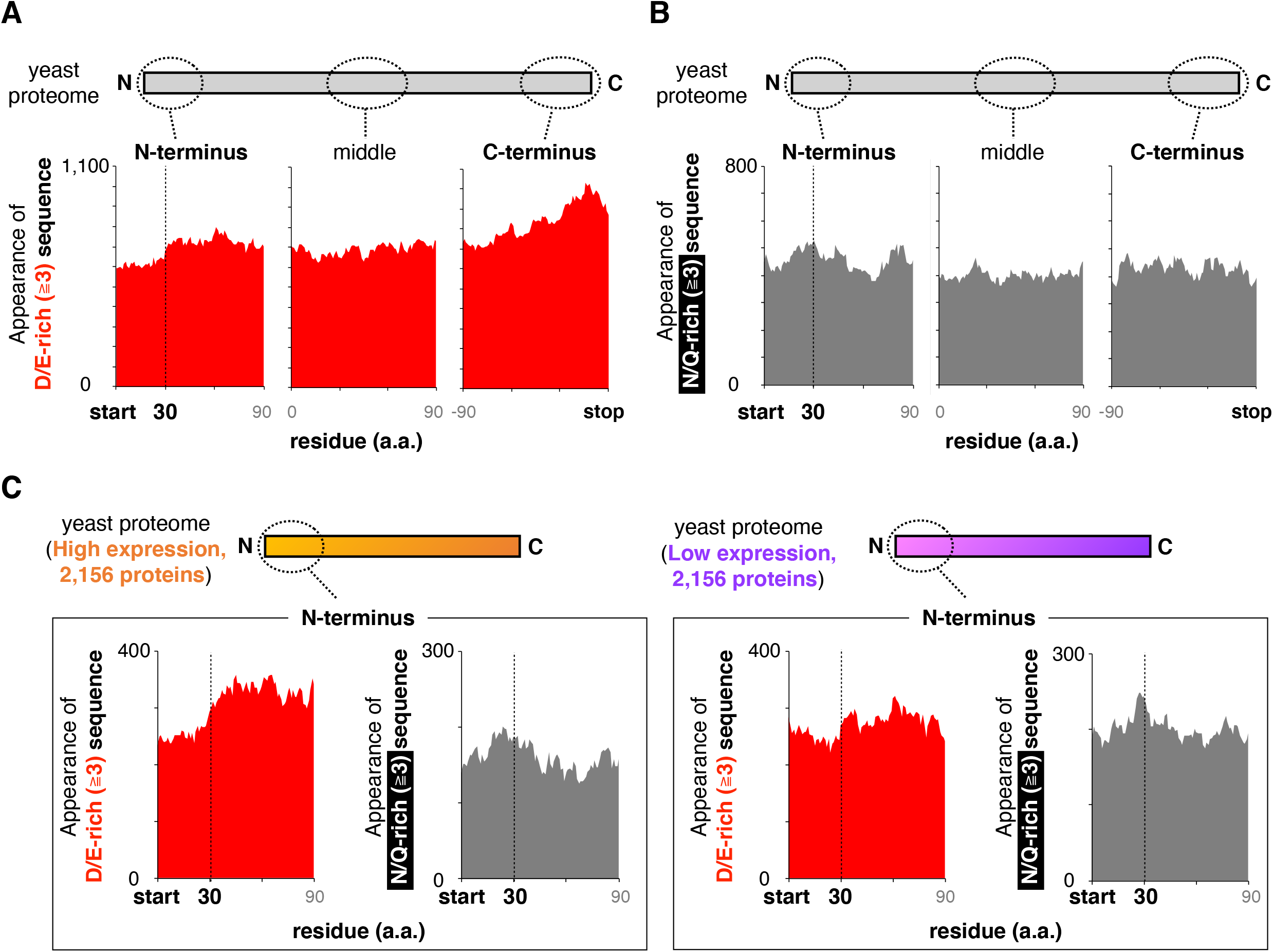
Yeast proteome avoids positioning D/E-rich sequences in the N-terminal regions. (**A and B**) Amino acid composition of the yeast proteome. The vertical axis represents the number of proteins harboring ≥3 D/E (**A**) or N/Q (**B**) in a window. N-terminus, C-terminus, and middle region are defined as N-terminal 100 amino acids excluding the first methionine, C-terminal 100 amino acids, and middle 100 amino acids of the 5,540 yeast proteins that have lengths greater than 130 amino acids. (**C**) Amino acid composition of the yeast proteome, depending on the protein abundance. The abundance data were obtained from the literature (Kulak *et al.*, 2014).

Further analyses using protein abundance data (Kulak *et al*, 2014) revealed that the bias for the disfavored D/E residues at the N-terminal regions was mainly attributed to abundant proteins (**Fig. 6C**). This tendency is reasonable, since the translation of abundant proteins harboring N-terminal D/E-rich sequences would be more problematic or wasteful. In addition, we observed this biased negative-charge amino acid distribution in the human proteome (**Fig. S7A**), consistent with our results using the HsPURE system. Collectively, the N-terminal D/E-rich peptide-dependent premature termination, which is an inherent translation defect, affects the amino acid distribution in a variety of proteomes.

## Discussion

We previously discovered a novel noncanonical ribosome behavior, IRD, in *E. coli* (Chadani *et al*., 2017), but it was not obvious that IRD could be applied to a eukaryotic translation system. Based on the following overlapping features revealed in this study, we conclude that IRD also occurs in eukaryotes. First, nascent polypeptide chains with consecutive D/E residues at the N-terminal regions eventually led to the premature termination of translation elongation. Second, the translation of such D/E-rich amino acid sequences produced abortive peptidyl-tRNAs that were sensitive to Pth2, indicating that the integrity of the translating ribosome is impaired or the peptidyl-tRNAs are released probably due to the destabilization of the ribosome. Third, the stabilization of the translating ribosomes by increasing the Mg^2+^ concentration or the tunnel-occupying nascent polypeptide (Chadani *et al*., 2017, 2021) alleviates the premature termination, reflecting that the stability of the ribosome is associated with the premature termination. Fourth, the premature termination took place in a human-based reconstituted cell-free translation system, indicating that only the essential translation factors are sufficient for the premature termination. Finally, the premature termination was stochastic during the translation of sequences enriched in negative-charge residues, since the MS analysis identified various aborted products including 6 to 10D in the GFP_5_-10D sequence. Since the translation elongation mechanism is generally well conserved among all kingdoms of life, the conservation of this noncanonical ribosome dynamics is not surprising. In particular, the ribosome tunnel is well conserved in a region close to the PTC (Dao Duck *et al*, 2019). We thus conclude that IRD is a conserved property of the ribosomes from *E. coli* to eukaryotes.

There are notable differences in IRD between *E. coli* and eukaryotes. The eukaryotic ribosomes are more robust to continuing the translation elongation of the D/E-runs. In *E. coli*, the stronger IRD motifs located in the internal regions of the ORF induce the translation abortion in a context-dependent manner. In contrast, our analysis showed that IRD-mediated translation abortion in eukaryotes only occurs when the D/E-rich sequences are located at the N-terminal regions. Indeed, there are many internal consecutive negatively charged residues in eukaryotic ORFs. The bioinformatic survey revealed that longer (≥10) D/E-runs appear more frequently in eukaryotes than bacteria (**Fig. S7B**). In budding yeast, dozens of genes encode more than 20 consecutive D/E residues in the middle of their ORFs. The most extreme one is Vhs3p (Ruiz *et al*, 2004, 2009), which includes 56 D/E runs starting from the 606th amino acid. The continuous translation after the long D/E runs was confirmed by the GWIPS-viz browser (Michel *et al*, 2013), a public web tool to visualize the ribosome distribution.

A possible explanation for the presence of such long D/E-rich sequences could be an extension of the protein functions using the regions in eukaryotes. The Gene Ontology (GO) analysis revealed that the human genes that are classified as being localized at the nucleus, chromosomes, and nucleolus, or binding to nucleic acids (DNA, RNA, histone, and chromatin) are enriched in D/E-runs (**Fig. S7C**), as pointed out in earlier studies (Karlin, 1995; Karlin *et al*, 2002; Lobanov *et al*, 2016). This suggests that the D/E-rich sequences are important for the acquisition of nuclear functions in eukaryotes. Domains including D/E-rich sequences can be regarded as “low (sequence) complexity” domains, which have recently received increasing attention as mediators to regulate liquid-liquid phase separation in the cell (Kato & McKnight, 2016; Alberti & Hyman, 2021). Indeed, the negatively charged Nephrin intracellular domain forms phase-separated nuclear bodies (Pak *et al*, 2016). Other studies also indicated that “hyperacidic” sequences are utilized as interacting domains or interaction regulators for various proteins (He *et al*, 1996, 1997; Zhu & Karlin, 1996; Ruiz *et al*, 2004, 2009; Zaharias *et al*, 2021). In addition, essential yeast proteins, such as the RNA binding protein Nab3p (Wilson *et al*, 1994; Fasken *et al*, 2015), the transcription elongation regulator Spt5p (Lindstrom *et al*, 2003; Schneider *et al*, 2006), and the 40S subunit biogenesis protein Kri1p (Sasaki *et al*, 2000; Hoareau-Aveilla *et al*, 2012) contain 23, 19 and 15 D/E runs from the 114th, 148th and 52nd residues within the 802, 1,063 and 591 aa length polypeptides, respectively. Collectively, stronger IRD counteraction would be reinforced to ensure the robust continuation of translation elongation for more complex eukaryotic proteomes.

Eukaryotic ribosomes are strengthened by several mechanisms, including eukaryote-specific intersubunit bridges such as eL19 and eL24 in the peripheral regions (Ben-Shem *et al*, 2011; Kisly *et al*, 2016). These features would make the eukaryotic ribosome association more robust to translate the D/E-rich sequences, as compared to the association of prokaryotic ribosomes. In addition, a recent comparative analysis of exit tunnel structures revealed that the tunnel in eukaryotes has a narrower part and a second constriction site (Dao Duc *et al*., 2019), suggesting a stronger interaction between the nascent chain and the interior of the tunnel. Our recent analysis in *E. coli* revealed the nascent chain “length”-dependent IRD counteraction at the early stage of translation, and we proposed that the nascent peptide chain in the exit tunnel has a built-in ability to ensure elongation continuity by serving as a “bridge” with the interior of the tunnel (Chadani *et al*., 2021). Eukaryotic translation systems also show frequent IRD at the N-terminal regions and the “length”-dependent IRD-counteraction, consistent with the properties of the bacterial IRD. Therefore, we assume that the narrower eukaryotic tunnel would provide a better chance of interactions and be a prerequisite to synthesize the “high-risk” sequences.

Recent studies revealed that eIF5A (Hyp2p) alleviates stalling on many motifs, including Asp-rich motifs such as DDG, besides the previously known proline stretches (Pelechano & Alepuz, 2017; Schuller *et al*, 2017), indicating that the negatively charged residues during elongation could also be problematic. An *in vivo* reporter analysis using the *hyp2* temperature-sensitive strain showed unaltered IRD-induced premature termination in the eIF5A-impaired cells, demonstrating that eIF5A is unlikely to rescue the defective ribosome in the IRD state, as EF-P, the bacterial homolog of eIF5A, is not associated with IRD in *E. coli* (Chadani *et al*., 2017). It is currently unclear why the DDG motif and consecutive D/E residues behave differently. Further studies, including detailed structural analyses, will be necessary to elucidate the differences between the two phenomena.

We found that only Pth2p is involved in the cleavage of polypeptidyl-tRNAs derived from IRD in yeast cells. *S. cerevisiae* possesses two peptidyl-tRNA hydrolases, Pth1p and Pth2p, orthologous to those in bacteria and archaea, respectively (Rosas-Sandoval *et al*, 2002). Previous comprehensive analyses annotated both Pths as localized to mitochondria (Sickmann *et al*, 2003; Huh *et al*, 2003). However, Ishii *et al* reported an interaction between Pth2p and the cytosolic Rad23-Dsk2 proteins that shuttle ubiquitinated proteins to the proteasome (Ishii *et al*, 2006). Thus we assume that Pth2p exists “on” the cytosolic side of the mitochondrial membrane and participates in not only the regulation of ubiquitinated protein degradation but also the clearance of the abortive peptidyl-tRNAs. In addition, the Pth2p-dependent inhibition of the substrate recruitment to the proteasome by Rad23-Dsk2 raises the interesting possibility that the IRD-derived peptides resolved by Pth2p could escape from degradation.

We developed an MS-based method to identify the endogenous peptidyl-tRNAs that are specific to cells lacking Pth2p. The properties of the identified peptidyl-tRNAs strongly suggested that IRD produces the peptidyl-tRNAs that are cleaved by Pth2p in wild-type cells. Although we identified ~130 *pth2*Δ-specific peptidyl-tRNAs by this method, we note that the scale of such drop-off products is probably underestimated. First, the peptidyl-tRNA species concentrated by the phenol-extraction would be biased and contain some nonspecific contaminants. Indeed, more than 80% of the peptidyl-tRNAs identified from the *pth2*Δ cells overlap with those from the wild-type cells. Second, the comprehensive identification of the entire peptidyl-tRNAs is hampered by technical limitations of MS, such as low ionization of the peptides and insufficient sensitivity. Despite these caveats, the method developed here clearly unveiled the prematurely terminated peptidyl-tRNA species in an IRD-dependent manner, which are hidden in wild-type cells.

In conclusion, our results definitively demonstrated that both prokaryotic and eukaryotic ribosomes encounter some difficulties in synthesizing consecutive negatively charged amino acid sequences. In terms of the continuation of elongation, IRD might be regarded as a defect in translation, but *E. coli* conversely harnesses this weakness of the translational machinery as an environmental sensor (Chadani *et al*., 2017). Although the physiological relevance of IRD in eukaryotes remains enigmatic, it is possible that IRD could be utilized for cellular functions such as gene expression regulation or expansion of polypeptide repertories. The bioinformatics analysis revealed that proteomes avoid D/E-rich sequences at the N-terminal position. This tendency is consistent with the previous reports for eukaryotes (Duc & Song, 2018; Tuller *et al*, 2011) and prokaryotes (Chadani *et al*., 2021). Furthermore, this is more obvious for genes with higher expression levels. Although this is reasonable for efficient protein synthesis, it should be noted that many other eukaryotic genes maintain the “risky” sequences that could induce translation abortion. These might reflect the demands for IRD to regulate cellular functions. Further studies are required to reveal the biological significance of IRD in eukaryotes.

## Author Contributions

Y.I., Y.C., T.N., and K.M. performed experiments; Y.I., Y.C., T.N., A.Y., K.M., H.I. and H.T. conceived the study, designed experiments and analyzed the results; Y.C. and H.T. supervised the entire project; Y.I., Y.C. and H.T. wrote the manuscript.

## Acknowledgments

We thank Kosuke Ito for providing Pth2; Takashi Kanamori for providing *E. coli* Pth; Motonori Ota for valuable discussions; Eri Uemura for technical support; and the Bio-support Center at Tokyo Tech for DNA sequencing. This work was supported by MEXT Grants-in-Aid for Scientific Research (Grant Numbers JP26116002, JP18H03984, JP20H05925 to HT, 17K15062, 19K16038 to YC) and a grant from the Ohsumi Frontier Science Foundation to YC.

## Materials and Methods

### Construction of yeast strains and plasmids

Yeast strains, plasmids, and oligonucleotides used in this study are listed in Tables S1, S2, and S3, respectively.

For gene deletions, a cassette containing only a selection marker was PCR amplified as described (Güldener *et al*, 1996, 2002), and introduced into the BY4741 background strain. All *Saccharomyces cerevisiae* strains and PCR-amplified DNA fragments used for gene disruption in this study are summarized in Table S1.

Plasmids were constructed using standard cloning procedures and Gibson assembly (Gibson, 2011). Detailed schemes are summarized in Table S2, and the sequences of constructed plasmids are available in the Mendeley repository.

### Yeast cultures

Yeast were cultivated at 30 °C in either liquid yeast extract–peptone–dextrose (YPD)-rich medium, in synthetic dextrose, galactose or raffinose based on casamino acids (SD/CA, SG/CA or SR/CA) minimal medium (6.7 g/L yeast nitrogen base without ammonium sulfate, with 5.0 g/L casamino acids and 2% glucose), or in synthetic dextrose, galactose or raffinose based on dropout (SD/DO, SG/DO or SR/DO) minimal medium (6.7 g/L yeast nitrogen base without ammonium sulfate with a dropout solution for each amino acid). Yeasts harboring plasmids for luciferase assays were grown in SD or SG lacking uracil; and yeasts harboring plasmids for the overexpression of yeast Pth1p, Pth2p or *E. coli* Pth were grown in SD or SG lacking leucine.

### Dual-luciferase reporter assay

Dual-luciferase assays were performed with the dual-luciferase reporter assay system (Promega Corp., Madison, WI). All reagents were prepared according to the manufacturer’s instructions. *S. cerevisiae* strains were grown to exponential phase in SR/CA medium. The cells were harvested by centrifugation and resuspended in SG/CA. After induction for 4 hours, a 10 μL portion of the culture was transferred to 100 μL of 1× passive lysis buffer. After allowing lysis to progress for 10 sec, a 10 μL aliquot was used for luminescence measurements with a Varioskan LUX Multimode Microplate Reader (Thermo Fisher Scientific). The following steps were performed on 96-well plates. A 50 μL portion of the firefly luciferase reagent (LARΠ) was added to the plate, with a 10 sec equilibration time and measurement of luminescence with a 10 sec integration time, followed by the addition of 50 μL of the *Renilla* luciferase reagent and firefly luciferase quenching (Stop & Glo), a 10 sec equilibration time, and measurement of luminescence with a 10 sec integration time. The data are represented as the ratio of Firefly to *Renilla* luciferase activity (Fluc activity / Rluc activity).

### Spot assay

*S. cerevisiae* strains were grown to exponential phase in appropriate media. The cultures were then serially diluted (10^7^, 10^6^, 10^5^, 10^4^ and 10^3^ cells/mL), and 4 μL portions of the diluted cell suspensions were spotted on appropriate plates and incubated at 30 °C for 2-3 days.

### RNA isolation for peptidyl tRNA analysis

*S. cerevisiae* strains were grown to exponential phase in SR/CA medium. The cells were harvested by centrifugation and resuspended in SG/CA. Total RNA was isolated using the TriPURE isolation reagent (Sigma-Aldrich) according to the manufacturer’s instructions.

### *In vitro* peptidyl-tRNA hydrolysis

The RNA sample was divided into four portions. Two were resuspended in HEPES-KOH (pH 7.6) and incubated with 1 μM of Pth2 at 30 °C for 20 min or 1 μM of Pth from *E. coli* at 37 °C for 20 min, and the others were resuspended in 100 mM Tris (pH 11.0) followed by heating at 80 °C for 20 min.

### Northern blot

RNA samples separated by 11% WIDE Range Gel SDS-PAGE were transferred onto a Hybond-N^+^ membrane (GE Healthcare) and hybridized with biotinylated oligonucleotide(s) (IDT) complementary to the tRNAs shown below with the probe sequences: tRNA^Asp^: CCGCGACGGGGAATTGAACCCCGATCTG, tRNA^Asn^: CCCCAGTGAGGGTTGAACTCACGATCTT, 5SrRNA ACCCACTACACTACTCGGTCAGGCTCTTAC. Hybridization experiments were performed using a NorthernMax kit (Ambion) and a Chemiluminescent Nucleic Acid Detection Module (Thermo Scientific) according to the manufacturer’s instructions. Images were visualized and analyzed by an LAS4000 LuminoImager (GE Healthcare).

### Sample preparation for LC-MS/MS analysis

For the quantification of the luciferase peptides, *S. cerevisiae* strains were grown to exponential phase in SR/CA medium. The cells were harvested by centrifugation and resuspended in SG/CA. Total protein was precipitated by adding an equal volume of 10% TCA followed by centrifugation (15,000 × g, 5 min, 4 °C). The precipitate was washed with ice-cooled acetone and centrifuged (15,000 × g, 5 min, 4 °C), and the supernatant was removed by aspiration. The resulting precipitant was solubilized by PTS buffer, consisting of 12 mM sodium deoxycholate (SDC), 12 mM sodium lauryl sulfate (SLS), and 100 mM Tris-HCl (pH 9.0). Protein concentrations were evaluated by using a Pierce BCA Protein Assay Kit (Thermo Scientific).

For the identification of the peptides from the peptidyl-tRNAs, RNA samples isolated by the TriPURE reagent and isopropanol precipitation were resuspended in PTS buffer, consisting of 12 mM sodium deoxycholate (SDC), 12 mM sodium lauryl sulfate (SLS), and 100 mM Tris (pH ~11). The solution was then incubated at 80 °C for 20 min to hydrolyze the peptidyl-tRNA. After the hydrolysis, PTS buffer consisting of 12 mM sodium deoxycholate (SDC), 12 mM sodium lauryl sulfate (SLS), and 100 mM Tris-HCl (pH 6.8) was added to adjust the pH to 9.

The obtained protein/peptide solution was reduced with 10 mM DTT for 30 min at room temperature, and the resultant proteins/peptides were alkylated with 50 mM iodoacetamide for 30 min at room temperature in the dark. After a 4-fold dilution with 50 mM ammonium bicarbonate, the proteins/peptides were digested with endoproteinase Lys-C (Wako Pure Chemical) at room temperature for 3 hours. Proteins/peptides were further digested with trypsin (Trypsin Gold, Promega) at 37 °C overnight. The amounts of added Lys-C or trypsin were 1 μg per 100 μg total protein or 1 μg per 50 μg total protein, respectively. After the digestion, SDC and SLS were removed by ethyl acetate extraction at low pH (0.5% trifluoroacetic acid). The samples were then desalted with a desalting column (GL-Tip SDB, GL Sciences) and dissolved in a 2% acetonitrile and 0.1% trifluoroacetic acid solution before the LC-MS/MS measurement.

### LC-MS/MS analysis

LC-MS/MS measurements were performed with a nanoLC-ESI-MS/MS system composed of a quadrupole-orbitrap hybrid mass spectrometer (Q-Exactive; Thermo Fisher Scientific) equipped with a nanospray ion source and a nano HPLC system (Easy-nLC 1000; Thermo Fisher Scientific). The trap column used for the nano HPLC was a 2 cm × 75 μm capillary column packed with 3 μm C18-silica particles (Thermo Fisher Scientific) and the separation column was a 12.5 cm × 75 μm capillary column packed with 3 μm C18-silica particles (Nikkyo Technos Co., Ltd.). The flow rate of the nano HPLC was 300 nL/min. The separation was conducted using a 10-40% linear acetonitrile gradient for 30 min for the identification of IRD candidates or 70 min for the quantification of luciferase peptides in the presence of 0.1% formic acid. The LC-MS/MS data were acquired in the data-dependent acquisition mode controlled by Xcalibur 4.0 (Thermo Fisher Scientific). The settings of the data-dependent acquisition were as follows: the resolutions were 70,000 for the full MS scan and 17,500 for the MS2 scan; the AGC targets were 3.0E6 for the full MS scan and 5.0E5 for the MS2 scan; the maximum IT was 60 ms for both the full MS and MS2 scans; the scan range was 310–1,500 m/z for the full MS scan, and the top 10 signals were selected for the MS2 scan per one full MS scan.

For the analysis of Firefly and *Renilla* luciferases, the Skyline software was used for the quantification of their MS1 intensities (MacLean *et al*, 2010). In this analysis, every ten peptides with high reproducibility were selected and used for comparison.

For the identification of the peptidyl-tRNAs, the MS/MS spectra were searched against all *S. cerevisiae* protein sequences obtained from the UniProt database, by using the MaxQuant software (ver. 1.6.9.0)(Cox & Mann, 2008) with the “Semi-specific free C-terminus” setting in the “Digestion” window. The identified peptides were defined based on the information from the “Identification type” column: the peptides detected at least once among the three measurements were defined as identified.

### *In vitro* translation and product analysis

The coupled transcription-translation reaction was performed using the HsPURE system (Machida *et al*., 2014) in the presence of ^35^S-methionine, at 32 °C for 90 min. The coupled transcription-translation reaction using the *E. coli* PURE system was performed by using PUREfrex (v1.0) as described previously (Chadani *et al*., 2017, 2021). Template DNAs for *in vitro* transcription-translation reactions of the mRNA encoding CrPV IRES were amplified by PCR, using PM0771(GGCCTAATACGACTCACTATAGGGAAAAAGC) and PM0773(GTTATTGCTCAGCGGTTAAGAGTTTTCACTGCATACGACG). In the case of the plasmids carrying HCV IRES, the template DNA was amplified by PCR using PM0511 and the GAL1_seq-primer (GCATAACCACTTTAACTAATACTTTCA). The template DNA for the *E. coli* PURE system was amplified by primer 1 (GGCCTAATACGACTCACTATAGGAGAAATCATAAAAAATTTATTTGCTTTGTGAG CGG) and primer 3 (AGTCAGTCACGATGAATTCCCCTAGCTTGG).

The reaction mixture was treated with 200 μM of puromycin for 5 min at 37 °C. Subsequently, the reaction mixture was divided into two portions, and one was incubated with 1 μM of Pth2p at 30 °C for 20 min. The reaction was stopped by dilution into an excessive volume of 5 % TCA. After standing on ice for at least 10 min, the samples were centrifuged for 3 min at 4 °C, and the supernatant was removed by aspiration. The precipitates were then vortexed with 0.9 mL of acetone, centrifuged again, and dissolved in SDS sample buffer (62.5 mM Tris-HCl, pH 6.8, 2% SDS, 10% glycerol, 50 mM DTT) that had been treated with RNasecure (Ambion). Finally, the sample was divided into two portions. One was incubated with 50 mg/mL of RNase A (Promega) at 37 °C for 30 min, and separated by the WIDE range SDS-PAGE system (Nacalai Tesque). The translation continuation index (TC index) was calculated by the following formula: {completed chain (10D)} / {completed chain (10D) + peptidyl-tRNA (10D)}.

### Amino acid composition analysis and enrichment analysis with Gene Ontology

The amino acid composition analysis was conducted with in-house R scripts (with R.app ver. 3.1.2). We defined the N-terminus, C-terminus, MID, and random regions as the N-terminal 100 amino acids excluding the first methionine, C-terminal 100 amino acids, and middle 100 amino acids of the 5,446 yeast proteins with lengths greater than 130 amino acids, and used these sequences to calculate the amino acid composition. Amino acid enrichment was calculated using a ten-residue moving window, and we counted proteins harboring more target amino acids than the determined number in a window. The same analysis was conducted for other organisms, as shown in Figure S7. All amino acid sequences were obtained from the UniProt database.

The enrichment analysis with Gene Ontology annotation for human proteome was conducted with the PANTHER web-tool (http://www.pantherdb.org/)(Thomas *et al*, 2003).

## Notes

### Competing Interest Statement

The authors have declared no competing interest.

## References

Alberti S & Hyman AA (2021) Biomolecular condensates at the nexus of cellular stress, protein aggregation disease and ageing. Nat Rev Mol Cell Bio 22: 196–213

Ben-Shem A, Loubresse NG de, Melnikov S, Jenner L, Yusupova G & Yusupov M (2011) The Structure of the Eukaryotic Ribosome at 3.0 Å Resolution. Science 334: 1524–1529

Bhushan S, Hoffmann T, Seidelt B, Frauenfeld J, Mielke T, Berninghausen O, Wilson DN & Beckmann R (2011) SecM-Stalled Ribosomes Adopt an Altered Geometry at the Peptidyl Transferase Center. PLoS Biol 9:e1000581

Bhushan S, Meyer H, Starosta AL, Becker T, Mielke T, Berninghausen O, Sattler M, Wilson DN & Beckmann R (2010) Structural Basis for Translational Stalling by Human Cytomegalovirus and Fungal Arginine Attenuator Peptide. Mol Cell 40: 138–146

Brandman O, Stewart-Ornstein J, Wong D, Larson A, Williams CC, Li G-W, Zhou S, King D, Shen PS, Weibezahn J, et al (2012) A Ribosome-Bound Quality Control Complex Triggers Degradation of Nascent Peptides and Signals Translation Stress. Cell 151: 1042–1054

Buskirk AR & Green R (2017) Ribosome pausing, arrest and rescue in bacteria and eukaryotes. Philos Trans R Soc Lond B Biol Sci 372: 20160183

Chadani Y, Niwa T, Chiba S, Taguchi H & Ito K (2016) Integrated in vivo and in vitro nascent chain profiling reveals widespread translational pausing. Proc National Acad Sci 113: E829–E838

Chadani Y, Niwa T, Izumi T, Sugata N, Nagao A, Suzuki T, Chiba S, Ito K & Taguchi H (2017) Intrinsic Ribosome Destabilization Underlies Translation and Provides an Organism with a Strategy of Environmental Sensing. Mol Cell 68: 528–539.e5

Chadani Y, Sugata N, Niwa T, Ito Y, Iwasaki S & Taguchi H (2021) Nascent polypeptide within the exit tunnel stabilizes the ribosome to counteract risky translation. EMBO J 40: e108299

Charneski CA & Hurst LD (2013) Positively charged residues are the major determinants of ribosomal velocity. PLoS Biol 11: e1001508

Choi J, Grosely R, Prabhakar A, Lapointe CP, Wang J & Puglisi JD (2015) How Messenger RNA and Nascent Chain Sequences Regulate Translation Elongation. Annu Rev Biochem 87: 421–449

Chomczynski P & Sacchi N (1987) Single-step method of RNA isolation by acid guanidinium thiocyanate-phenol-chloroform extraction. Anal Biochem 162: 156–159

Cox J & Mann M (2008) MaxQuant enables high peptide identification rates, individualized p.p.b.-range mass accuracies and proteome-wide protein quantification. Nat Biotechnol 26: 1367–1372

Cui Y, Hagan KW, Zhang S & Peltz SW (1995) Identification and characterization of genes that are required for the accelerated degradation of mRNAs containing a premature translational termination codon. Gene Dev 9: 423–436

Dao Duc K, Batra SS, Bhattacharya N, D. Cate JH & Song YS (2019) Differences in the path to exit the ribosome across the three domains of life. Nucleic Acids Res: gkz106–

Doma MK & Parker R (2006) Endonucleolytic cleavage of eukaryotic mRNAs with stalls in translation elongation. Nature 440: 561–564

Duc KD & Song YS (2018) The impact of ribosomal interference, codon usage, and exit tunnel interactions on translation elongation rate variation. Plos Genet 14: e1007166

Fasken MB, Laribee RN & Corbett AH (2015) Nab3 Facilitates the Function of the TRAMP Complex in RNA Processing via Recruitment of Rrp6 Independent of Nrd1. Plos Genet 11: e1005044

Gibson DG (2011) Chapter 15 - Enzymatic Assembly of Overlapping DNA Fragments 1st ed. Elsevier Inc.

Gutierrez E, Shin B-S, Woolstenhulme CJ, Kim J-R, Saini P, Buskirk AR & Dever TE (2013) eIF5A Promotes Translation of Polyproline Motifs. Mol Cell 51: 35–45

He F, Brown AH & Jacobson A (1996) Interaction between Nmd2p and Upf1p is required for activity but not for dominant-negative inhibition of the nonsense-mediated mRNA decay pathway in yeast. RNA 2: 153–70

He F, Brown AH & Jacobson A (1997) Upf1p, Nmd2p, and Upf3p are interacting components of the yeast nonsense-mediated mRNA decay pathway. Mol Cell Biol 17: 1580–1594

He F & Jacobson A (1995) Identification of a novel component of the nonsense-mediated mRNA decay pathway by use of an interacting protein screen. Gene Dev 9: 437–454

Heurgué-Hamard V, Mora L, Guarneros G & Buckingham RH (1996) The growth defect in Escherichia coli deficient in peptidyl-tRNA hydrolase is due to starvation for Lys-tRNA(Lys). EMBO J 15: 2826–2833

Hoareau-Aveilla C, Fayet-Lebaron E, Jády BE, Henras AK & Kiss T (2012) Utp23p is required for dissociation of snR30 small nucleolar RNP from preribosomal particles. Nucleic Acids Res 40: 3641–3652

Huh W-K, Falvo JV, Gerke LC, Carroll AS, Howson RW, Weissman JS & O’Shea EK (2003) Global analysis of protein localization in budding yeast. Nature 425: 686–691

Inada T (2017) The Ribosome as a Platform for mRNA and Nascent Polypeptide Quality Control. Trends Biochem Sci 42: 5–15

Ishii T, Funakoshi M & Kobayashi H (2006) Yeast Pth2 is a UBL domain-binding protein that participates in the ubiquitin–proteasome pathway. EMBO J 25: 5492–5503

Ito K, Chadani Y, Nakamori K, Chiba S, Akiyama Y & Abo T (2011) Nascentome Analysis Uncovers Futile Protein Synthesis in Escherichia coli. PLoS ONE 6: e28413

Ito K & Chiba S (2013) Arrest Peptides: Cis-Acting Modulators of Translation. Annu Rev Biochem 82: 171–202

Karlin S (1995) Statistical significance of sequence patterns in proteins. Curr Opin Struc Biol 5: 360–371

Karlin S, Brocchieri L, Bergman A, Mrázek J & Gentles AJ (2002) Amino acid runs in eukaryotic proteomes and disease associations. Proc Natl Acad Sci USA 99: 333–338

Kato M & McKnight SL (2016) A Solid-State Conceptualization of Information Transfer from Gene to Message to Protein. Annu Rev Biochem 87: 1–40

Kim JH, Lee S-R, Li L-H, Park H-J, Park J-H, Lee KY, Kim M-K, Shin BA & Choi S-Y (2011) High Cleavage Efficiency of a 2A Peptide Derived from Porcine Teschovirus-1 in Human Cell Lines, Zebrafish and Mice. Plos One 6: e18556

Kisly I, Gulay SP, Mäeorg U, Dinman JD, Remme J & Tamm T (2016) The Functional Role of eL19 and eB12 Intersubunit Bridge in the Eukaryotic Ribosome. J Mol Biol 428: 2203–2216

Kulak NA, Pichler G, Paron I, Nagaraj N & Mann M (2014) Minimal, encapsulated proteomic-sample processing applied to copy-number estimation in eukaryotic cells. Nat Methods 11: 319–324

Kuroha K, Zinoviev A, Hellen CUT & Pestova TV (2018) Release of Ubiquitinated and Non-ubiquitinated Nascent Chains from Stalled Mammalian Ribosomal Complexes by ANKZF1 and Ptrh1. Mol Cell 72: 286–302.e8

Li Z, Vizeacoumar FJ, Bahr S, Li J, Warringer J, Vizeacoumar FS, Min R, VanderSluis B, Bellay J, DeVit M, et al (2011) Systematic exploration of essential yeast gene function with temperature-sensitive mutants. Nat Biotechnol 29: 361–367

Lindstrom DL, Squazzo SL, Muster N, Burckin TA, Wachter KC, Emigh CA, McCleery JA, Yates JR & Hartzog GA (2003) Dual Roles for Spt5 in Pre-mRNA Processing and Transcription Elongation Revealed by Identification of Spt5-Associated Proteins. Mol Cell Biol 23: 1368–1378

Lobanov MYu, Klus P, Sokolovsky IV, Tartaglia GG & Galzitskaya OV (2016) Non-random distribution of homo-repeats: links with biological functions and human diseases. Sci Rep 6: 26941

Lu J & Deutsch C (2008) Electrostatics in the Ribosomal Tunnel Modulate Chain Elongation Rates. J Mol Biol 384: 73–86

Machida K, Mikami S, Masutani M, Mishima K, Kobayashi T & Imataka H (2014) A translation system reconstituted with human factors proves that processing of encephalomyocarditis virus proteins 2A and 2B occurs in the elongation phase of translation without eukaryotic release factors. J Biol Chem 289: 31960–31971

MacLean B, Tomazela DM, Shulman N, Chambers M, Finney GL, Frewen B, Kern R, Tabb DL, Liebler DC & MacCoss MJ (2010) Skyline: an open source document editor for creating and analyzing targeted proteomics experiments. Bioinformatics 26: 966–968

Matsuo Y, Ikeuchi K, Saeki Y, Iwasaki S, Schmidt C, Udagawa T, Sato F, Tsuchiya H, Becker T, Tanaka K, et al (2017) Ubiquitination of stalled ribosome triggers ribosome-associated quality control. Nat Commun 8: 1–13

Menez J, Buckingham RH, Zamaroczy M de & Campelli CK (2002) Peptidyl-tRNA hydrolase in Bacillus subtilis, encoded by spoVC, is essential to vegetative growth, whereas the homologous enzyme in Saccharomyces cerevisiae is dispensable. Mol Microbiol 45: 123–129

Menninger JR (1979) Accumulation of peptidyl tRNA is lethal to Escherichia coli. J Bacteriol 137: 694–696

Michel AM, Fox G, Kiran AM, Bo CD, O’Connor PBF, Heaphy SM, Mullan JPA, Donohue CA, Higgins DG & Baranov PV (2013) GWIPS-viz: development of a ribo-seq genome browser. Nucleic Acids Res 42: D859–D864

Pak CW, Kosno M, Holehouse AS, Padrick SB, Mittal A, Ali R, Yunus AA, Liu DR, Pappu RV & Rosen MK (2016) Sequence Determinants of Intracellular Phase Separation by Complex Coacervation of a Disordered Protein. Mol Cell 63: 72–85

Pelechano V & Alepuz P (2017) eIF5A facilitates translation termination globally and promotes the elongation of many non polyproline-specific tripeptide sequences. Nucleic Acids Res 45: 7326–7338

Rendón OZ, Fredrickson EK, Howard CJ, Vranken JV, Fogarty S, Tolley ND, Kalia R, Osuna BA, Shen PS, Hill CP, et al (2018) Vms1p is a release factor for the ribosome-associated quality control complex. Nat Commun 9: 1–9

Ron EZ, Kohler RE & Davis BD (1968) Magnesium ion dependence of free and polysomal ribosomes from Escherichia coli. J Mol Biol 36: 83–89

Rosas-Sandoval G, Ambrogelly A, Rinehart J, Wei D, Cruz-Vera LR, Graham DE, Stetter KO, Guarneros G & Söll D (2002) Orthologs of a novel archaeal and of the bacterial peptidyl–tRNA hydrolase are nonessential in yeast. Proc Natl Acad Sci USA 99: 16707–16712

Ruiz A, González A, Muñoz I, Serrano R, Abrie JA, Strauss E & Ariño J (2009) Moonlighting proteins Hal3 and Vhs3 form a heteromeric PPCDC with Ykl088w in yeast CoA biosynthesis. Nat Chem Biol 5: 920–928

Ruiz A, Muñoz I, Serrano R, González A, Simón E & Ariño J (2004) Functional characterization of the Saccharomyces cerevisiae VHS3 gene: a regulatory subunit of the Ppz1 protein phosphatase with novel, phosphatase-unrelated functions. J Biol Chem 279: 34421–34430

Sasaki T, Toh-e A & Kikuchi Y (2000) Yeast Krr1p physically and functionally interacts with a novel essential Kri1p, and both proteins are required for 40S ribosome biogenesis in the nucleolus. Mol Cell Biol 20: 7971–7979

Schlessinger D, Mangiarotti G & Apirion D (1967) The formation and stabilization of 30S and 50S ribosome couples in Escherichia coli. Proc Natl Acad Sci USA 58: 1782–9

Schneider DA, French SL, Osheim YN, Bailey AO, Vu L, Dodd J, Yates JR, Beyer AL & Nomura M (2006) RNA polymerase II elongation factors Spt4p and Spt5p play roles in transcription elongation by RNA polymerase I and rRNA processing. Proc Natl Acad Sci USA 103: 12707–12712

Schuller AP, Wu CC-C, Dever TE, Buskirk AR & Green R (2017) eIF5A Functions Globally in Translation Elongation and Termination. Mol Cell 66: 194–205.e5

Seidelt B, Innis CA, Wilson DN, Gartmann M, Armache J-P, Villa E, Trabuco LG, Becker T, Mielke T, Schulten K, et al (2009) Structural Insight into Nascent Polypeptide Chain–Mediated Translational Stalling. Science 326: 1412–1415

Shanmuganathan V, Schiller N, Magoulopoulou A, Cheng J, Braunger K, Cymer F, Berninghausen O, Beatrix B, Kohno K, Heijne G von, et al (2019) Structural and mutational analysis of the ribosome-arresting human XBP1u. eLife 8: e46267

Sickmann A, Reinders J, Wagner Y, Joppich C, Zahedi R, Meyer HE, Schönfisch B, Perschil I, Chacinska A, Guiard B, et al (2003) The proteome of Saccharomyces cerevisiae mitochondria. Proc Natl Acad Sci USA 100:13207–13212

Thomas PD, Campbell MJ, Kejariwal A, Mi H, Karlak B, Daverman R, Diemer K, Muruganujan A & Narechania A (2003) PANTHER: A Library of Protein Families and Subfamilies Indexed by Function. Genome Res 13: 2129–2141

Tuller T, Veksler-Lublinsky I, Gazit N, Kupiec M, Ruppin E & Ziv-Ukelson M (2011) Composite effects of gene determinants on the translation speed and density of ribosomes. Genome Biol 12: R110–R110

Verma R, Reichermeier KM, Burroughs AM, Oania RS, Reitsma JM, Aravind L & Deshaies RJ (2018) Vms1/Ankzf1 peptidyl-tRNA hydrolase releases nascent chains from stalled ribosomes. Nature 557: 446–451

Wen J-D, Lancaster L, Hodges C, Zeri A-C, Yoshimura SH, Noller HF, Bustamante C & Tinoco I (2008) Following translation by single ribosomes one codon at a time. Nature 452: 598–603

Wilson DN & Beckmann R (2011) The ribosomal tunnel as a functional environment for nascent polypeptide folding and translational stalling. Curr Opin Struc Biol 21: 274–282

Wilson SM, Datar KV, Paddy MR, Swedlow JR & Swanson MS (1994) Characterization of nuclear polyadenylated RNA-binding proteins in Saccharomyces cerevisiae. J Cell Biol 127: 1173–1184

Wolin SL & Walter P (1988) Ribosome pausing and stacking during translation of a eukaryotic mRNA. EMBO J 7: 3559–3569

Yanofsky C (1981) Attenuation in the control of expression of bacterial operons. Nature 289: 751–758

Zaharias S, Zhang Z, Davis K, Fargason T, Cashman D, Yu T & Zhang J (2021) Intrinsically disordered electronegative clusters improve stability and binding specificity of RNA-binding proteins. J Biol Chem 297: 100945

Zhu ZY & Karlin S (1996) Clusters of charged residues in protein three-dimensional structures. Proc Natl Acad Sci USA 93: 8350–8355

