## Supplemental Figures for "Nascent peptide-induced translation discontinuation in eukaryotes impacts biased amino acid usage in proteomes"

**Supplemental Figures S1-S7**

**Figure S1**: Detailed analysis of translation abortion caused by negatively charged amino acids. Related to Fig. 1.

**Figure S2**: Features of IRD in eukaryotes evaluated by a reconstituted cell free translation system, the HsPURE system. Related to Fig. 2.

**Figure S3**: Additional experiments on the function of Pth2. Related to Fig. 3.

**Figure S4**: Mapping of the LC-MS/MS-identified peptides derived from the peptidyl-tRNAs. Related to Fig. 4.

**Figure S5**: Additional experiments on IRD during the translation of endogenous genes detected in the *pth2*Δ strain by LC-MS/MS analysis. Related to Fig. 5.

**Figure S6**: Data analysis of yeast proteome sequences. Related to Fig. 6.

**Figure S7**: Distribution of the D/E-enriched sequences among all kingdoms of life. Related to Fig. 6.

**Supplemental Tables S1-S3**

**Table S1**: Strains used in this study.

**Table S2**: Plasmids used in this study.

**Table S3**: Oligonucleotides used in this study.

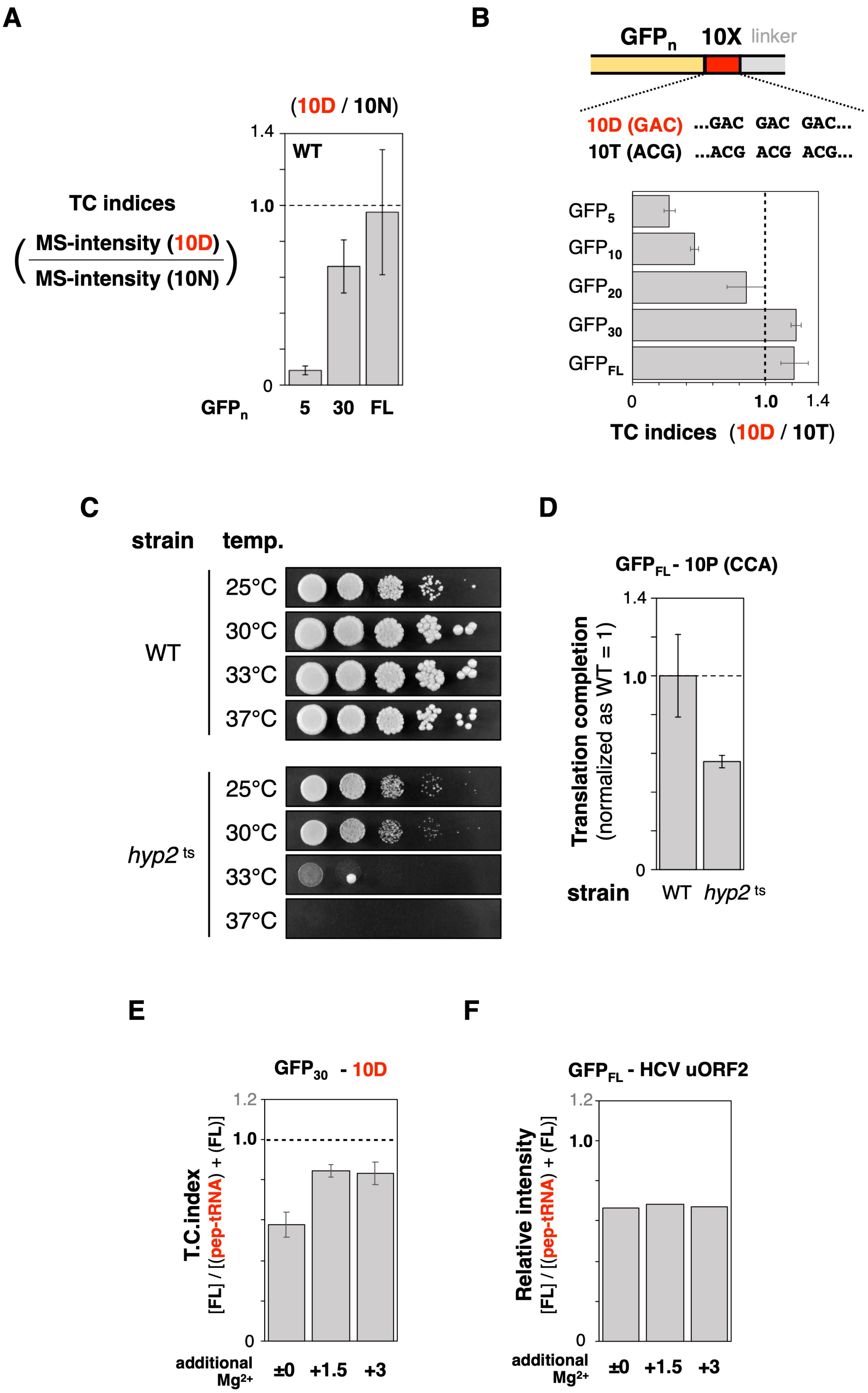
A
(10D / 10N)
WT
TC indices
MS-intensity (10D)
MS-intensity (10N)
GFPn
5
30
FL
B
GFPn
10X
linker
10D (GAC) ...GAC GAC GAC...
10T (ACG) ...ACG ACG ACG...
GFP5
GFP10
GFP20
GFP30
GFPFL
TC indices (10D / 10T)
C
strain
temp.
WT
hyp2 ts
25°C
30°C
33°C
37°C
D
GFPFL - 10P (CCA)
Translation completion
(normalized as WT = 1)
strain
WT
hyp2 ts
E
GFP30 - 10D
T.C.index
[FL] / [(pep-tRNA) + (FL)]
additional Mg2+
±0
+1.5
+3
F
GFPFL - HCV uORF2
Relative intensity
[FL] / [(pep-tRNA) + (FL)]
additional Mg2+
±0
+1.5
+3

**Figure S1**

**Detailed analysis of translation abortion caused by negatively charged amino acids. Related to Fig. 1.**

(**A**) The relative quantification of the luciferases by MS analysis. Ten peptides in each luciferase were detected with LC-MS/MS and their MS1 intensities were quantified with the Skyline software. Translation continuation (TC) indices were calculated as follows; first, the 10D/10N ratio was calculated for each peptide and the ratios between Fluc and Rluc were calculated for all the combinations of the ten peptides in each luciferase ($10\times10 = 100$ combinations). Then, the mean value of the 100 Fluc/Rluc ratios was depicted as a barplot. Error bars indicate standard deviations of 100 Fluc/Rluc ratios. (**B**) Translation attenuation depends on the negatively charged amino acid sequence but not the nucleotide sequence in the vicinity of the N-terminal region *in vivo*. The TC indices were calculated by comparing the relative Fluc activity between 10D (GAC) and 10T (ACG, -1 frameshift), as shown in **Fig. 1B**. (**C and D**) Confirmation of *hyp2*$^{ts}$ strain, TSA737. Wild-type (WT) and *hyp2*$^{ts}$ strains were spotted on the YPD plate at indicating temperature for 2 days. The *hyp2*$^{ts}$ strain shows a semi-lethal phenotype at 33 °C (**C**) and the inefficiency of a poly-proline (10P) translation at 33 °C (**D**). (**E**) Effect of $Mg^{2+}$ on the 10D sequence-dependent translation attenuation. The $GFP_{30}$-10D sequences were translated by the HsPURE system supplemented with additional $Mg^{2+}$ as indicated, and individual TC indices were calculated. (**F**) Effect of $Mg^{2+}$ on the hCMV uORF-induced translational stalling. The relative intensity of the peptidyl-tRNAs calculated from two-independent experiments was shown.

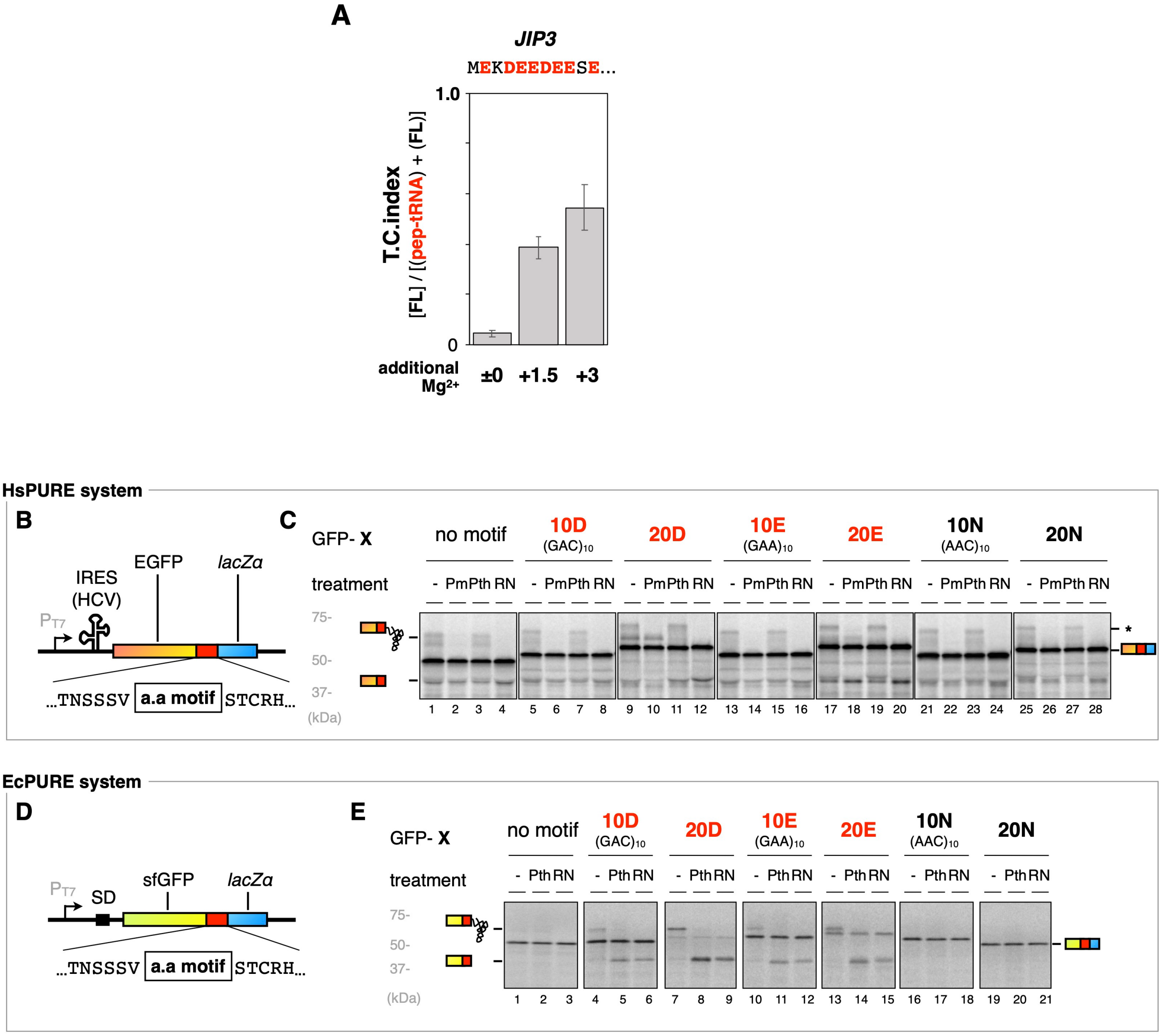


**Figure S2**

**Features of IRD in eukaryotes evaluated by a reconstituted cell free translation system, the HsPURE system.**

(**A**) An endogenous D/E-rich sequence attenuates translation elongation in an $Mg^{2+}$-dependent manner. The fusion of the N-terminal 10 a.a. of *JIP3* and a truncated Fluc (**Fig. 2D**) was translated by the HsPURE system supplemented with additional $Mg^{2+}$ as indicated, and individual TC indices were calculated, as shown in **Fig. 1G**.

(**B**) Schematic representation of the EGFP-10X model sequence for the HsPURE *in vitro* translation assay.

(**C**) The EGFP-10X genes were translated using the HsPURE system including $^{35}$S-methionine as shown in **Fig. 1E**. The products were treated with a peptidyl-tRNA hydrolase (yeast Pth2p: *Pth*) or puromycin (Pm) as indicated and separated by neutral pH SDS-PAGE with optional RNase A (*RN*) pretreatment. Radioactive bands were detected by a phosphorimager. Full-length product, peptidyl-tRNAs and the tRNA-cleaved polypeptides are indicated by schematic labels.

(**D**) Schematic representation of the sfGFP-10X model sequence for an *E. coli* factor-based reconstituted cell-free translation system (PUREfrex v1.0, EcPURE).

(**E**) The sfGFP-10X genes were translated using the EcPURE including $^{35}$S-methionine. The products were treated with a peptidyl-tRNA hydrolase (*E. coli* : *Pth*) as indicated and separated by neutral pH SDS-PAGE with optional RNase A (*RN*) pretreatment.

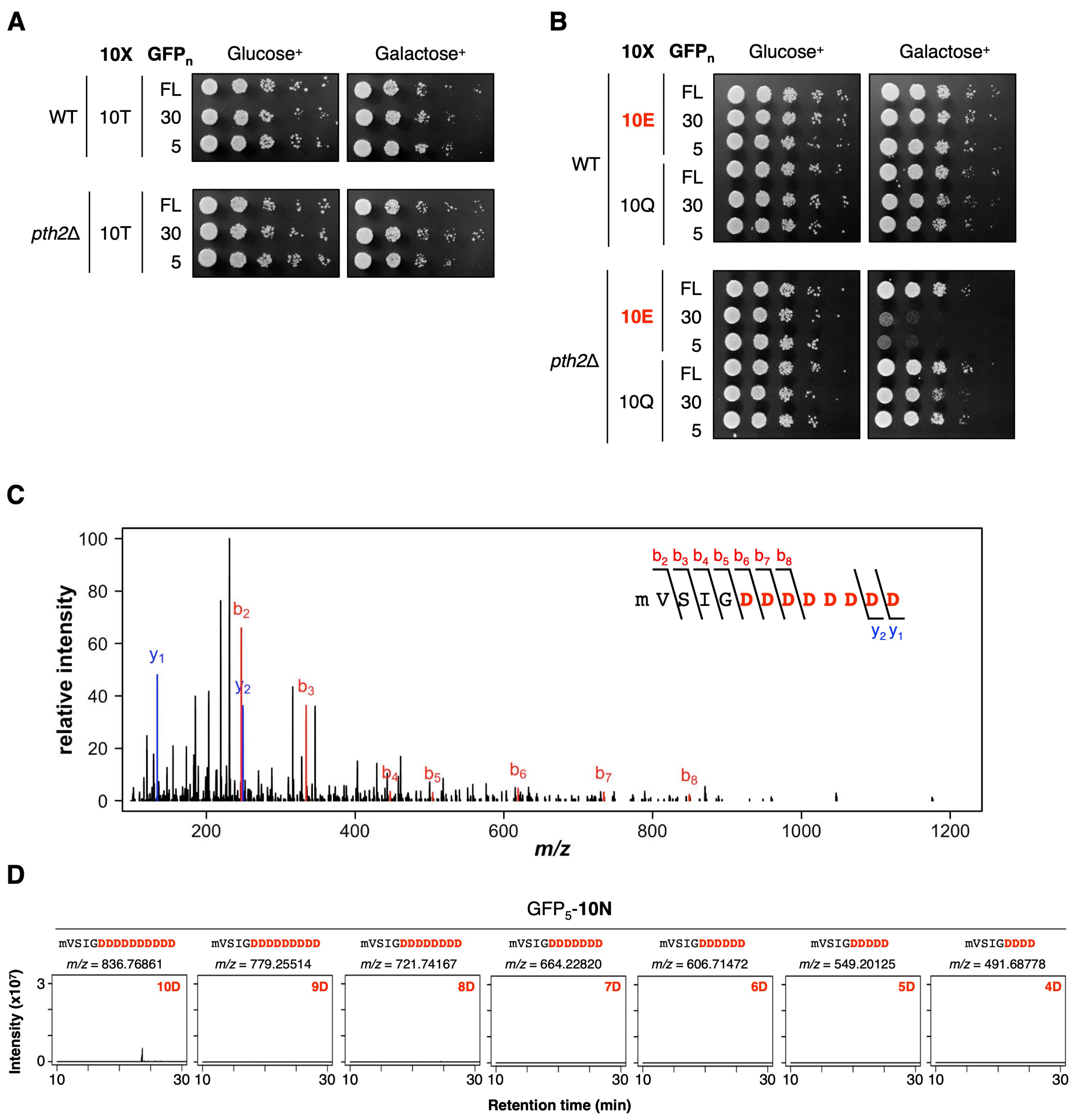


**Figure S3**

**Additional experiments on the function of Pth2. Related to Fig. 3.**

(**A**) Wild-type (WT) and *pth2*Δ strains harboring the plasmids for the expression of the $GFP_X$-10T shown in **Fig. S1B** were spotted on SD or SG plate lacking uracil at 30 °C for 2-3 days.

(**B**) Wild-type (WT) and *pth2*Δ strains harboring inducible plasmids shown in **Fig. 1B** (10E or 10Q series) were spotted on SD or SG plate lacking uracil at 30 °C for 2-3 days.

(**C**) The MS/MS spectrum of the $M_{ox}$VSIGDDDDDDDDDD peptide fragment expressed from $GFP_5$-10D. The peaks of the b- and y- ion fragments aare shown in red and blue, respectively.

(**D**) Scarce accumulation of the peptidyl-tRNAs, which were produced in the translation of $GFP_5$-10N, as a control for $GFP_5$-10D (**Fig. 3E**). Each panel represents extracted ion chromatograms for a monoisotopic ion of a $GFP_5$-(X)D peptide with a specific m/z value as indicated at the top.

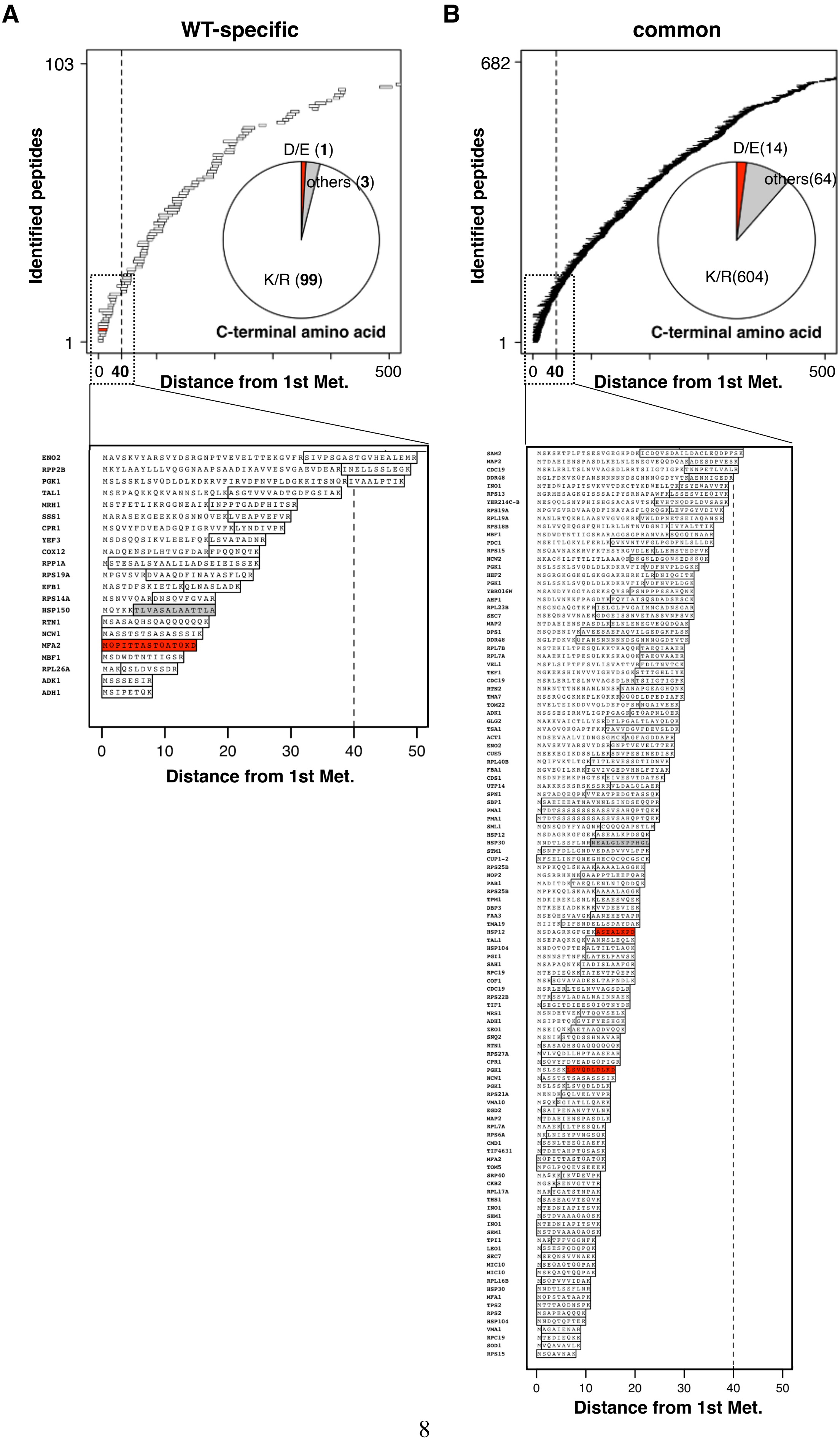
A
WT-specific
103
1
Identified peptides
D/E (1)
others (3)
K/R (99)
C-terminal amino acid
0
40
500
Distance from 1st Met.
0
10
20
30
40
50
Distance from 1st Met.
B
common
682
1
Identified peptides
D/E(14)
others(64)
K/R(604)
C-terminal amino acid
0
40
500
Distance from 1st Met.
0
10
20
30
40
50
Distance from 1st Met.

**Figure S4**

**Mapping of the LC-MS/MS-identified peptides derived from the peptidyl-tRNAs. Related to Fig. 4.**

(**A and B**) The mapping of identified peptides derived from the peptidyl-tRNAs, which are specific to wild-type (WT) (**A)** and are common in wild-type and *pth2*Δ strains (**B**). Map charts representing the localization of identified peptides on each protein, with the enlarged view presented below. The dotted lines indicate the position of 40 amino acids from the first methionine. The length of the 40 amino acids is roughly equivalent to that of the exit tunnel in the ribosome. *Insets*: Pie charts representing the amino acids at the C-termini of identified peptides (same as **Fig. 4E**, *red*: D/E; gray: others; white: K/R).

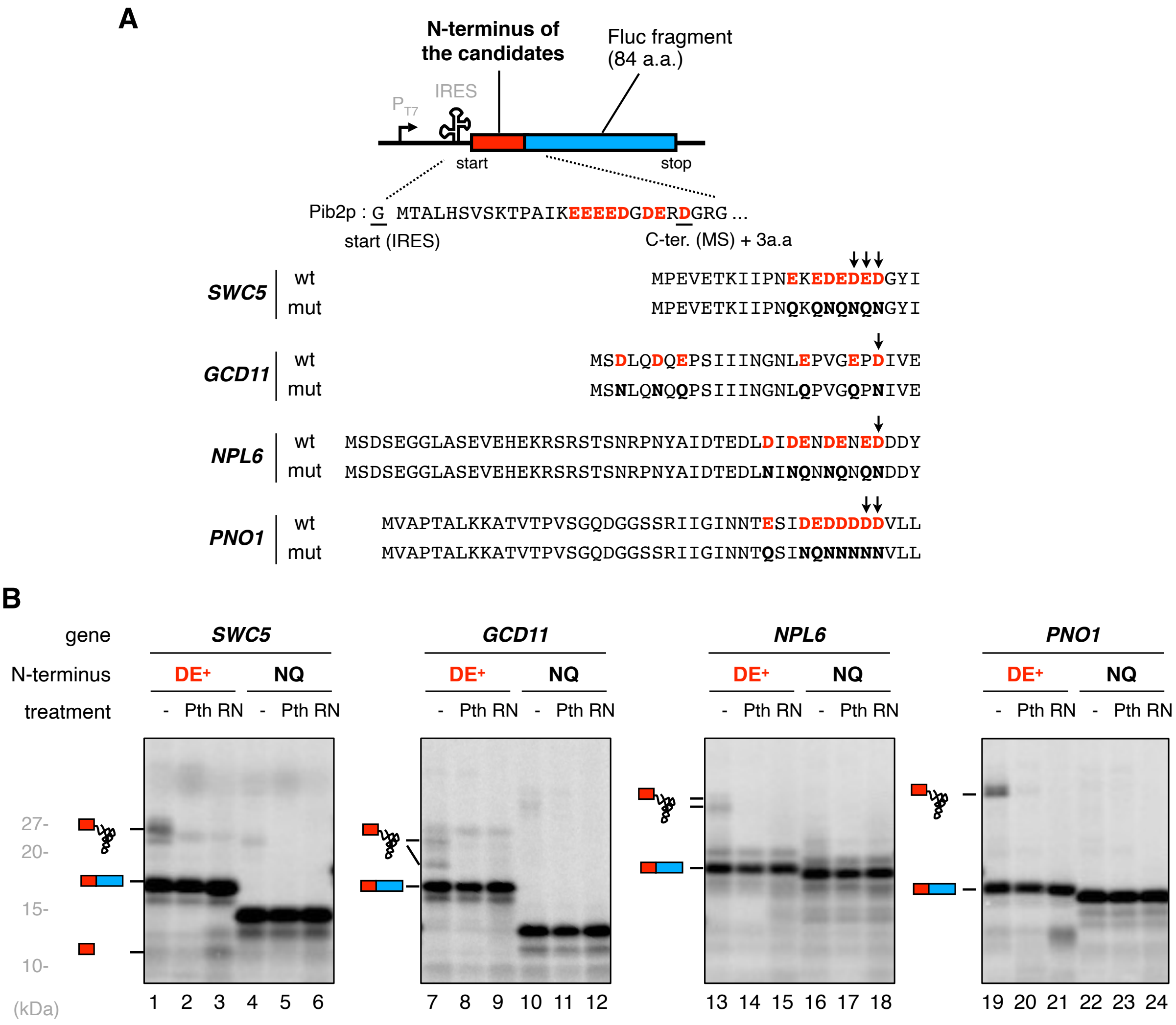


### Figure S5

**Additional experiments on IRD during the translation of endogenous genes detected in the *pth2*Δ strain by LC-MS/MS analysis. Related to Fig. 5.**

(**A**) Schematic representation of the sequences from endogenous genes for the HsPURE *in vitro* translation assay as described in **Fig. 5B**. (**B**) We expanded the analysis in **Fig. 5B** for *SWC5, GCD11*, *NPL6* and *PNO1*, and $^{35}$S-labeled *in vitro* translation products were separated by SDS-PAGE as described in **Fig. 1E**.

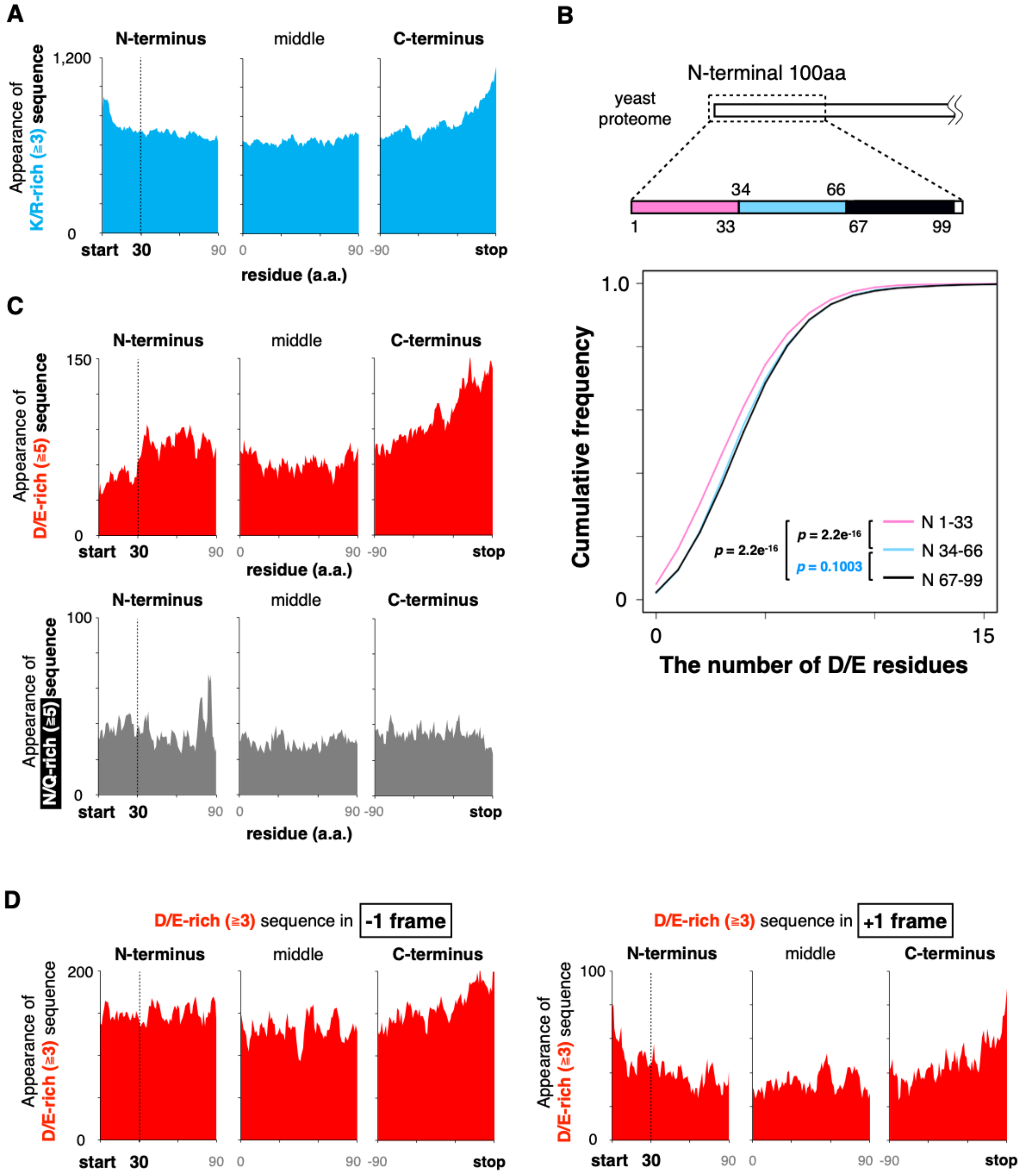


**Figure S6**

**Data analysis of yeast proteome sequences. Related to Fig. 6.**

(**A**) The composition of positively charged amino acids in the yeast proteome. The vertical axis represents the number of protein harboring ≥3 K/R in a window. N-terminus, C-terminus, and middle region are defined as N-terminal 100 amino acids excluding the first methionine, C-terminal 100 amino acids, and middle 100 amino acids of the yeast 5,540 yeast proteins that have lengths greater than 130 amino acids.

(**B**) Cumulative frequency of the D/E number within each one third part of N-terminal 100 amino acid residues of the yeast proteome. Wilcoxon rank sum test was used to compare the regions.

(**C**) The number of protein harboring ≥5 D/E (**upper**) or N/Q (**bottom**) in a window.

(**D**) Amino acids composition of the yeast proteome to confirm independency of a nucleic acid sequence bias. The vertical axis represents the number of proteins harboring ≥3 D/E in a -1 frameshift window (**left**) or a +1 frameshift window (**right**).

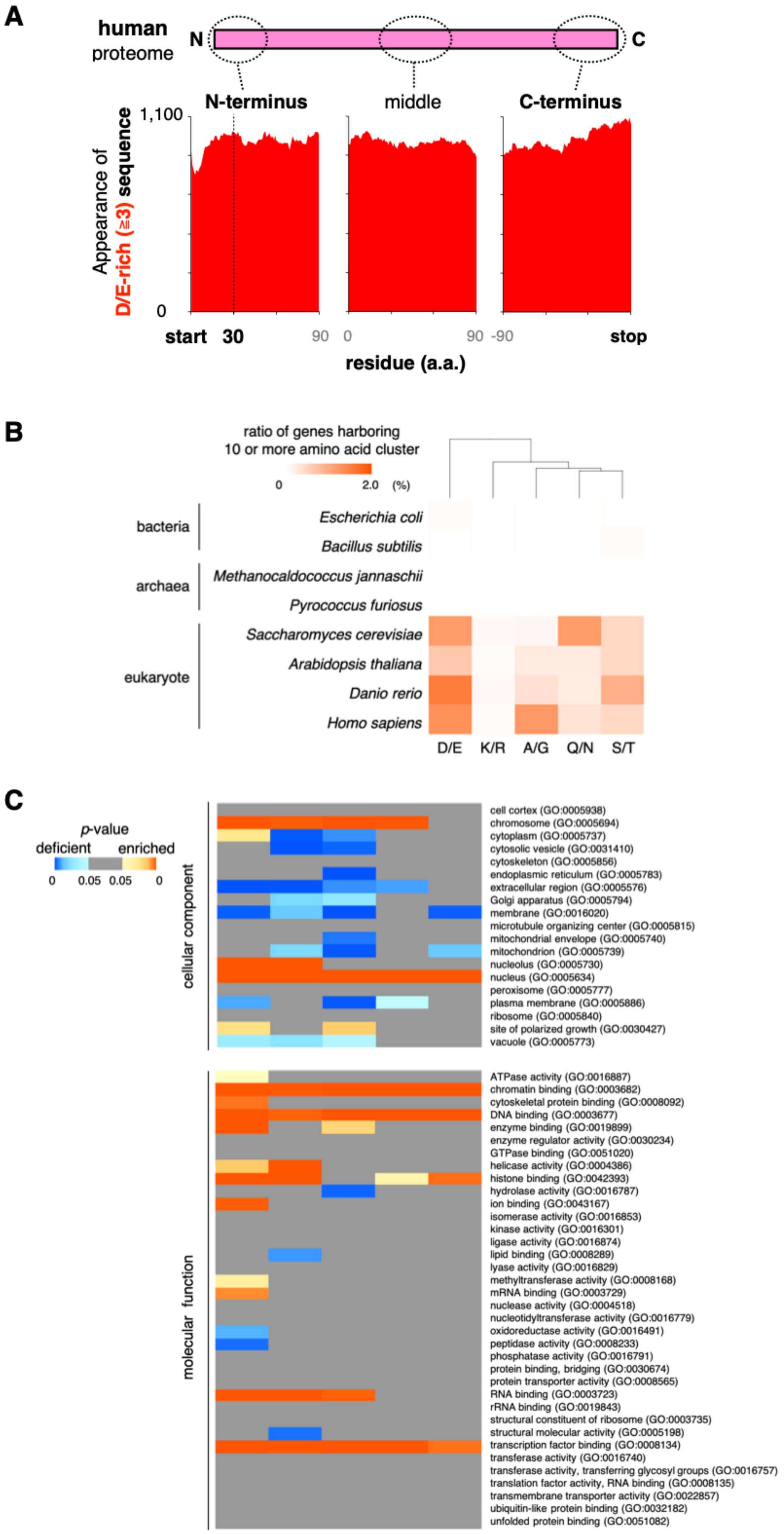
A
human
proteome
N
C
N-terminus
middle
C-terminus
1,100
0
Appearance of
D/E-rich (≥3) sequence
start
30
90
0
90
-90
stop
residue (a.a.)
B
ratio of genes harboring
10 or more amino acid cluster
0
2.0
(%)
bacteria
Escherichia coli
Bacillus subtilis
archaea
Methanocaldococcus jannaschii
Pyrococcus furiosus
eukaryote
Saccharomyces cerevisiae
Arabidopsis thaliana
Danio rerio
Homo sapiens
D/E
K/R
A/G
Q/N
S/T
C
p-value
deficient
enriched
0
0.05
0.05
0
cellular component
cell cortex (GO:0005938)
chromosome (GO:0005694)
cytoplasm (GO:0005737)
cytosolic vesicle (GO:0031410)
cytoskeleton (GO:0005856)
endoplasmic reticulum (GO:0005783)
extracellular region (GO:0005576)
Golgi apparatus (GO:0005794)
membrane (GO:0016020)
microtubule organizing center (GO:0005815)
mitochondrial envelope (GO:0005740)
mitochondrion (GO:0005739)
nucleolus (GO:0005730)
nucleus (GO:0005634)
peroxisome (GO:0005777)
plasma membrane (GO:0005886)
ribosome (GO:0005840)
site of polarized growth (GO:0030427)
vacuole (GO:0005773)
molecular function
ATPase activity (GO:0016887)
chromatin binding (GO:0003682)
cytoskeletal protein binding (GO:0008092)
DNA binding (GO:0003677)
enzyme binding (GO:0019899)
enzyme regulator activity (GO:0030234)
GTPase binding (GO:0051020)
helicase activity (GO:0004386)
histone binding (GO:0042393)
hydrolase activity (GO:0016787)
ion binding (GO:0043167)
isomerase activity (GO:0016853)
kinase activity (GO:0016301)
ligase activity (GO:0016874)
lipid binding (GO:0008289)
lyase activity (GO:0016829)
methyltransferase activity (GO:0008168)
mRNA binding (GO:0003729)
nuclease activity (GO:0004518)
nucleotidyltransferase activity (GO:0016779)
oxidoreductase activity (GO:0016491)
peptidase activity (GO:0008233)
phosphatase activity (GO:0016791)
protein binding, bridging (GO:0030674)
protein transporter activity (GO:0008565)
RNA binding (GO:0003723)
rRNA binding (GO:0019843)
structural constituent of ribosome (GO:0003735)
structural molecular activity (GO:0005198)
transcription factor binding (GO:0008134)
transferase activity (GO:0016740)
transferase activity, transferring glycosyl groups (GO:0016757)
translation factor activity, RNA binding (GO:0008135)
transmembrane transporter activity (GO:0022857)
ubiquitin-like protein binding (GO:0032182)
unfolded protein binding (GO:0051082)

**Figure S7**

**Distribution of the D/E-enriched sequences among all kingdoms of life. Related to Fig. 6.**

(**A**) Amino acids composition of the human proteome. The number of gene having three or more D/E in a window. The vertical axis represents the number of protein harboring ≥3 target amino acids in a window.

(**B**) Frequency of the genes containing ten consecutive amino acid sequences in bacteria, archaea, and eukaryotes. Frequency was indicated by the intensity of the orange color.

(**C**) Gene ontology enrichment analysis for the human ORFs containing the ten consecutive amino acid sequences. The enriched or deficient terms are indicated by warm or cold colors, respectively.
